# Convergent gene erosion in the chemical defensome of marine mammals

**DOI:** 10.64898/2026.05.21.726804

**Authors:** Bram Danneels, Diogo Oliveira, L. Filipe C. Castro, Odd André Karlsen, Raquel Ruivo, Anders Goksøyr

**Affiliations:** Computational Biology Unit, Department of Informatics, University of Bergen, Norway; Interdisciplinary Centre of Marine and Environmental Research, University of Porto, Portugal; Department of Biology, Faculty of Sciences, University of Porto, Portugal; Department of Biological Sciences, University of Bergen, Norway

## Abstract

To preserve homeostasis in the face of continual chemical insult, animals evolved dedicated molecular systems that detect, detoxify, and eliminate foreign compounds. Collectively, these enzymes, transporters, and regulatory pathways constitute the chemical defensome. In cetaceans, the loss of two key nuclear receptors (NR1I2/PXR and NR1I3/CAR) suggests a profound rearrangement of the chemical defense systems. Therefore, we investigated the gene inventory of the chemical defensome in Cetacea and two other major marine mammal lineages (Pinnipedia and Sirenia), using their closest terrestrial relatives to understand the extent and patterns of chemical defensome remodelling. We demonstrate large-scale gene loss in chemical defensome genes of cetaceans, as well as smaller scale gene loss in the other two marine mammal lineages, indicating possible convergent evolution. Gene loss occurred predominantly in phase I and phase II biotransformation enzymes, including CYPs, FMOs, SULTs, and GSTs. Many of the lost genes in cetaceans are known to be regulated by PXR and/or CAR, while genes lost in multiple marine mammal lineages are often not regulated by these transcription factors. We hypothesize that the transition to aquatic environments, often accompanied by corresponding changes in feeding habits, led to convergent loss of chemical defensome genes, and loss of PXR and CAR in cetaceans accelerated these losses. These findings reveal systematic erosion of chemical defense capabilities across marine mammal lineages, suggesting that adaptation to marine life involves trade-offs in detoxification capacity that may have significant implications for these species’ responses to increasing chemical pollution in present-day ocean environments.

## Introduction

Transcriptional regulators play central roles in coordinating gene expression across biological pathways. When such regulators are lost during evolution, an important question arises: what becomes of the genes and functions they control? This question is particularly relevant in the scope of the chemical defensome, an integrated network of genes involved in sensing, metabolizing, and eliminating harmful compounds (Goldstone et al., 2006). In response to chemical challenges, animals rely on these ancient endogenous detoxification systems that have been co-opted to mitigate the effects of novel contaminants (Croom, 2012). The chemical defensome can be broadly divided into four major categories: transcription factors, biotransformation pathways, transporters, and stress responses (Franco et al., 2025a; Goldstone et al., 2006; Xu et al., 2005). As regulators of gene expression, transcription factors coordinate a myriad of molecular responses to all sorts of aggression. One subgroup of these are the nuclear receptors (NRs), with the distinguishing feature of containing a ligand-binding domain that can modulate their activity upon ligand binding (Goksøyr, S.Ø, in prep). Some of these NRs, such as the PXR (pregnane X receptor) and CAR (constitutive androstane receptor), constitute the first line of defence against chemical threats, as they can detect foreign compounds and coordinate the expression of all the defensome machinery (Rakateli et al., 2023; Tolson and Wang, 2010; Xu et al., 2005). More downstream, biotransformation enzymes modify the foreign compounds to make them more hydrophilic, facilitating elimination or excretion from the cell and/or organism, transporters export altered chemicals out of the cells, while stress response genes such as antioxidants, heat-, and metal-responsive proteins protect cells from derivative products such as reactive oxygen species (Xu et al., 2005). Recently, the chemical defensome concept has been expanded to include avoidance behaviour, whereby organisms flee from contaminated areas as part of the *extended* chemical defensome (Franco et al., 2025a). Together, these systems form a multilayered defence network that determines an organism’s capacity to detect, process, and tolerate chemical stressors.

Since its original description in the sea urchin, the chemical defensome has been investigated in a range of other species including various invertebrates (De Marco et al., 2017; Goldstone, 2008; Roncalli et al., 2025; Shinzato et al., 2012; Yadetie et al., 2012), and fish (Eide et al., 2021; Franco et al., 2025b). Despite being critical for survival, these studies highlighted species-specific nuances. Thus, identifying the inventory of chemical defence genes is a key step in determining the capacity of a species to deal with xenobiotic and potentially harmful compounds. However, in spite of such growing interest, its composition and evolutionary dynamics remain poorly understood in marine mammals. Marine mammals, including whales and dolphins, are a particularly susceptible group, with high concentrations of contaminants detected in their tissues (Braune et al., 2005; El-Sharkawy et al., 2025; Jepson et al., 2016). Their long lifespans, high trophic positions, and lipid-rich tissues make them especially prone to the bioaccumulation and biomagnification of persistent organic pollutants and other toxic compounds (Reckendorf et al., 2023). Although their chemical defensome is not yet fully characterised, sparse reports suggest changes in specific chemical defensome genes. For instance, paraoxonase 1, a bloodstream enzyme that prevents lipid oxidation, was shown to be pseudogenized in marine mammals, decreasing defences against organophosphorus compounds (Meyer et al., 2018). In addition, cetaceans are known to possess fewer cytosolic glutathione S-transferase (GST) genes when compared to terrestrial mammals (Tian et al., 2019). It has also been suggested they have a lower number of genes coding for the cytochrome P450 monooxygenases (CYPs) and UDP-glucuronosyltransferases (UGTs) (Kim et al., 2016). Similarly, changes in CYPs, GSTs, UGTs, and other biotransformation gene families such as sulfotransferases (SULTs) have been described in pinnipeds (Kakehi et al., 2015; Kondo et al., 2023, 2022, 2017; Tian et al., 2019). Perhaps the most impactful change to the chemical defensome in marine mammals, is the loss of both PXR and CAR in cetaceans, whereas these nuclear receptors are retained in other marine mammal lineages such as pinnipeds, sirenians, and polar bears (Hecker et al., 2019; Lille-Langøy et al., 2015; Wagner et al., 2022). This contrast raises the possibility of lineage-specific evolutionary remodelling of xenobiotic sensing and metabolism in marine mammals. Given the increasing accumulation of environmental chemicals as a consequence of human activity, such historical remodelling could be posing serious challenges to wildlife conservation efforts, notably in the marine environment where anthropogenic chemical pollution is of particular concern (Chen et al., 2024; Noël and Brown, 2021).

In this work we use recently available high-quality reference genomes from large-scale genome sequencing projects, such as the Earth Biogenome Project and the Darwin Tree of Life, to provide a fine-scale characterization of the chemical defensome in cetaceans and other marine mammals (Lewin et al., 2018; Morin et al., 2025; The Darwin Tree of Life Project Consortium, 2022). Specifically, we aim to identify lineage-specific changes, and investigate whether cetaceans show signatures of defensome remodeling associated with the loss of these key xenosensors. We hypothesise that the loss of PXR and CAR in cetaceans has been accompanied by evolutionary shifts in other defensome components, reflecting either compensatory adaptation or altered constraints on xenobiotic response pathways. By providing a comparative genomic view of the chemical defensome in marine mammals, this study contributes to a broader understanding of how species adapt to chemically challenging environments and may help inform future assessments of pollutant vulnerability in marine megafauna.

## Results

### The chemical defensome in marine mammals and their relatives

We used orthology inference and functional predictions to identify the gene inventory of the chemical defensome in three marine mammal lineages (Cetacea, Pinnipedia, and Sirenia) and their closest relatives (terrestrial Artiodactyla, Carnivora, and Afrotheria). The number and proportion of chemical defensome genes are relatively stable within the different investigated lineages, with an average of 474 identified genes related to the chemical defensome (Table S1). Proportionally, the defensome takes up a similar percentage of the total predicted proteome in the different lineages, ranging from 2.25% in terrestrial carnivorans to 2.44% in terrestrial cetartiodactyls. However, pseudogene prediction based on alignment with human proteins predicted an average of 44 chemical defensome genes as pseudogenes, leading to a predicted functional chemical defensome of 430 genes on average. When only considering functional genes, the chemical defensome-related genes make up between 1.88% and 2.43% of the total predicted proteome. Terrestrial cetartiodactyls have the highest proportion of chemical defensome genes (2.24%), followed by terrestrial afrotheres (2.17%), and carnivorans (both terrestrial and aquatic; 2.08%). Cetaceans and the Florida manatee have the smallest proportions: 2.02% and 1.96% on average. The latter two lineages also have the highest proportion of predicted pseudogenes in their chemical defensomes: 13.53% and 15.66% of the identified chemical defensome genes are predicted to be pseudogenes in cetaceans and the Florida manatee, respectively. The proportion of predicted pseudogenes is significantly higher in these lineages than in their terrestrial relatives: 8.40% in terrestrial cetartiodactyls and 7.52% in terrestrial afrotheres. Interestingly, pinnipeds also show an increased proportion of predicted pseudogenes (9.85%) compared to their terrestrial relatives (7.35%), even though the size of the functional chemical defensome is similar in both lineages.

The chemical defensome can be broadly divided into four groups of genes: transcription factors, biotransformation genes, transporters, and stress-related genes. When looking at the distributions of these categories, we found that stress-responsive genes and biotransformation genes take up the largest part of the chemical defensome, followed by membrane transporters (Fig. 1). The stress-responsive genes mainly consist of heat-shock proteins, while antioxidants and metal-responsive genes are among the smallest groups of chemical defensome genes. While we did not observe any significant differences between marine mammals and their closest relatives in stress responses as a whole, we did observe a significantly higher number of antioxidant genes in pinnipeds compared to terrestrial carnivorans (31.78 vs. 27.53), and in cetaceans compared to terrestrial cetartiodactyls (27.19 vs. 24.69). The number of functional transcription factors is relatively stable among lineages, while we identified significantly fewer biotransformation genes (oxygenases, reductases, and transferases) and transporters in cetaceans compared to their terrestrial relatives (Fig. 1). In pinnipeds we only identified significantly fewer phase I biotransformation genes compared to their closest relatives, which is only due to a significantly lower number of oxygenases. While we could not identify any statistically significant differences between sirenians and the other afrotheres (due to having only one sirenian genome, the Florida manatee), we did observe lower numbers of oxygenases, phase II biotransformation genes, transporters, and heat-responsive proteins (Fig. 1).

**Figure 1:**
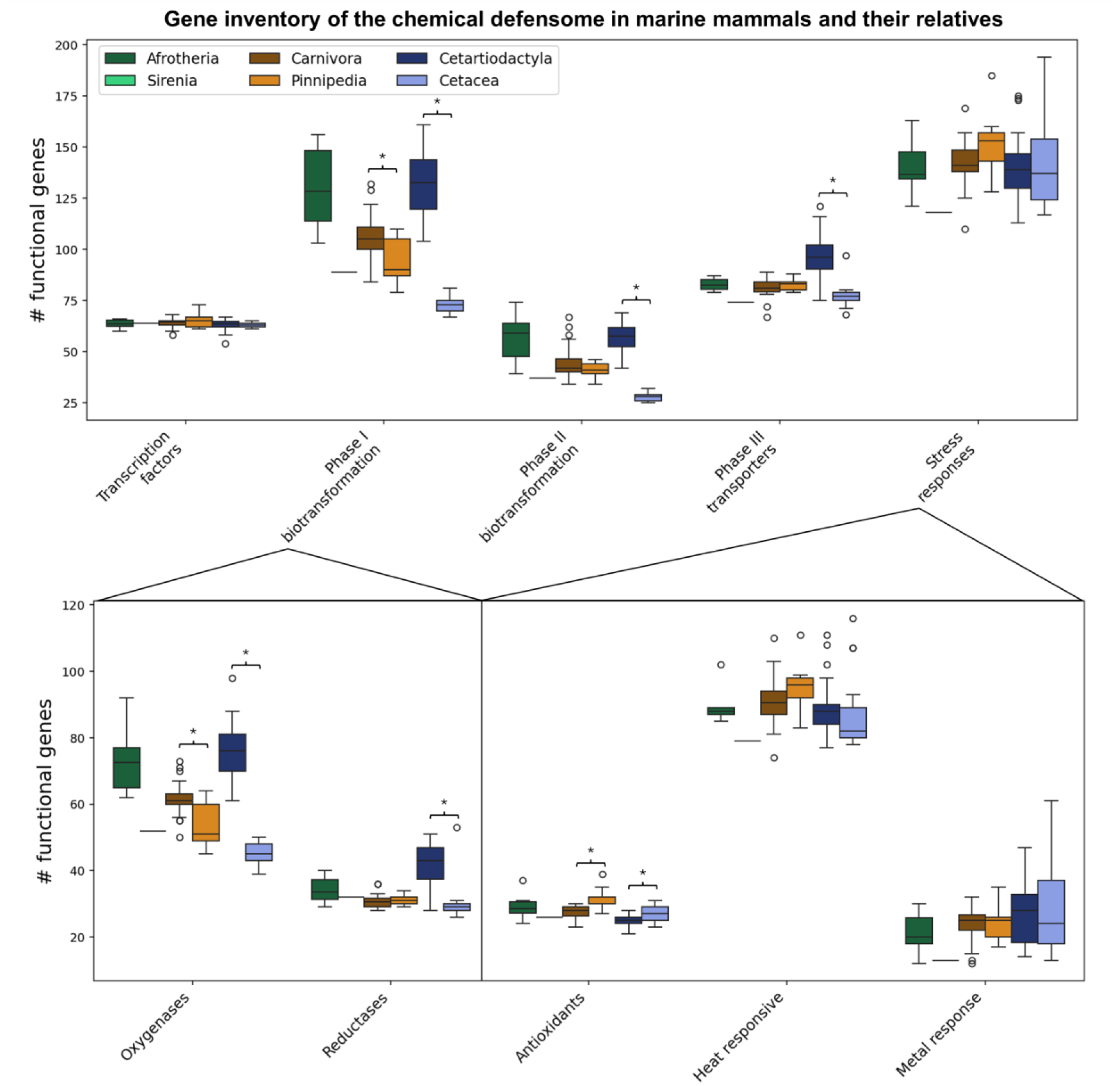
Functional gene counts in chemical defensome-related functional categories across target mammal lineages. Boxes represent medians and interquartile range. Circles represent outliers falling outside of 1.5x the interquartile range. Horizontal brackets with a star indicate statistically significant differences (p<0.05) in gene counts between marine lineages and the closest relative lineage, based on Mann-Whitney *U* tests (Crawford-Howell t-test in case of Sirenia vs. other Afrotheria).

### Transcription factors

As expected, transcription factors are generally well conserved among the investigated lineages (Fig. 2). However, we did identify lineage-specific differences in some transcription factor families. We confirmed the loss of both NR1I2 (PXR) and NR1I3 (CAR) in all investigated cetaceans, while being generally conserved in all other lineages. Although the PXR and CAR genes can still be detected in the cetacean genomes, they all showed multiple signs of pseudogenization. Another nuclear receptor, NR1H5 (FXR-β), is not found in the genomes of ruminants (Ruminantia) nor pigs (Suina), but was detected in all other investigated species, including other cetartiodactyls such as camels (Tylopoda), the hippopotamus, and cetaceans. NCOA3 (nuclear receptor coactivator 3) showed signs of pseudogenization in cetaceans and a select few other species, but further investigation of the protein-to-genome alignments revealed the impact of these alterations are likely minimal and could potentially still provide functional proteins. In addition, most carnivorans and a few cetartiodactyls have an extra NCOA5-like gene in their chemical defensome. Lastly, PER3 (period circadian protein 3) showed signs of pseudogenization in many different species, but is generally better conserved in marine species. For example, PER3 is retained in nearly all cetaceans, while it is pseudogenized in nearly all other cetartiodactyls. Similarly, it showed signs of possible pseudogenization in only three out of nine pinnipeds, while similar gene degradation was found in 27 out of the 34 other carnivorans.

**Figure 2:**
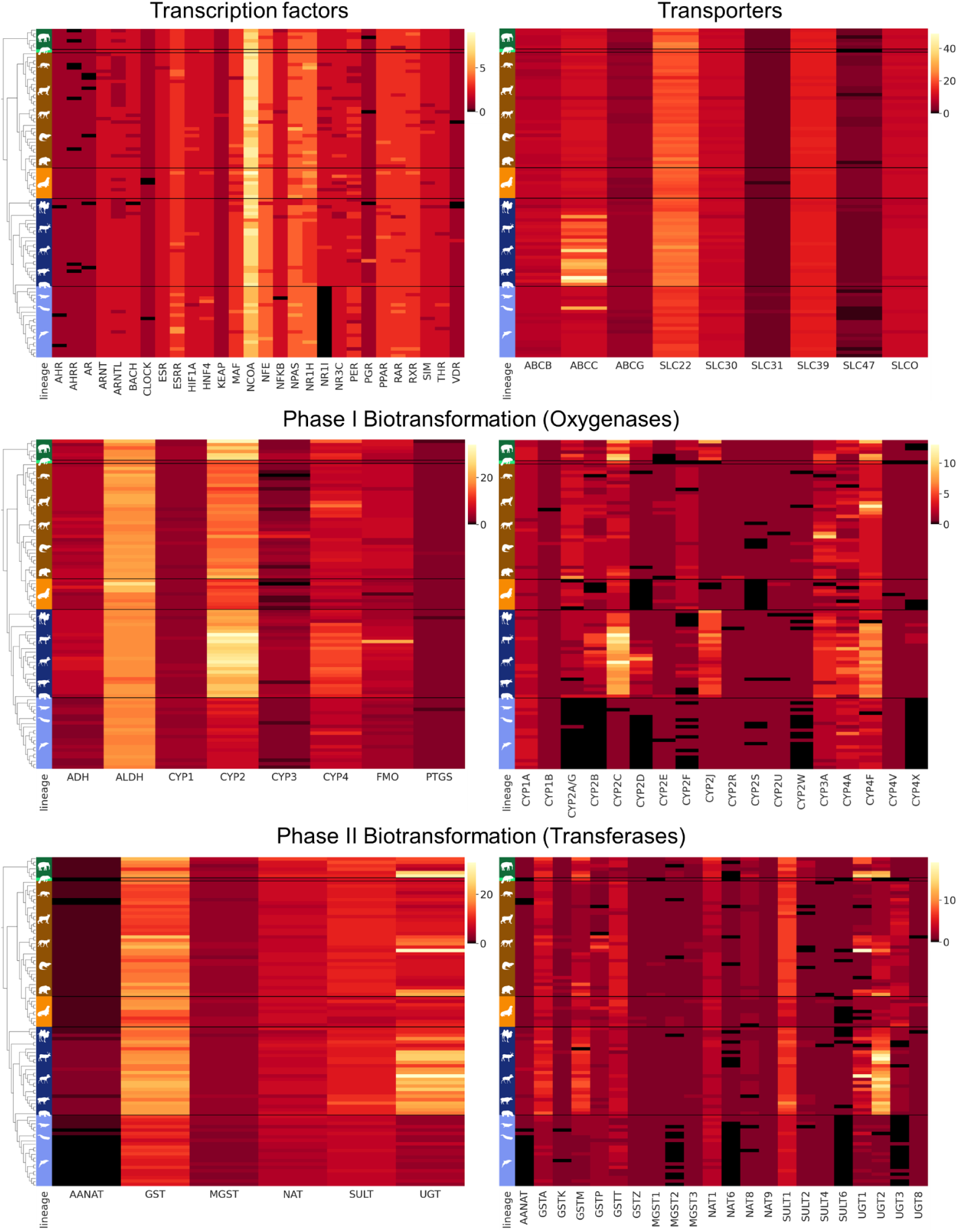
Gene count and distribution of transcription factors, oxygenases, and transferases of the chemical defensome across Cetartiodactyla, Carnivora, and Afrothera. Colours on the left correspond to the different investigated lineages. Dark green: Afrotheria (without Sirenia); light green: Sirenia; brown: Carnivora (without Pinnipedia); orange: Pinnipedia; dark blue: Cetartiodactyla (without Cetacea); light blue: Cetacea. Heatmap colours correspond to the number of detected functional genes in each gene family.

### Phase I biotransformation

A large part of the chemical defensome consists of biotransformation enzymes, which convert substances into more polar, water-soluble metabolites. Many of the genes involved in this process tend to organize in clusters, and for the most variable of them in phases I and II of biotransformation we constructed synteny maps with representative species for comparison. (Figs. 3 and S2-S6). We also checked liver gene expression within these most variable clusters in publicly available RNA-sequencing runs to further validate gene pseudogenization (Figs. S7-S11). The majority of enzymes involved in the first phase of biotransformation are CYPs, but other oxygenases (e.g. flavin-containing monooxygenases, or FMOs) and reductases (e.g. aldo-keto reductases, or AKRs) also take part in this process. We identified a relatively similar number of phase I reductases between lineages, except for the terrestrial cetartiodactyls (Fig. 1). In this group, we identified an increased number of AKRs, especially in the ruminants (Fig. S1). The expansion of AKRs in ruminants seems to be exclusively due to significant expansion of the AKR1C subfamily, with up to 17 copies in some species. In cetaceans, we observed pseudogenization of the reductase NQO1 (NAD(P)H dehydrogenase [quinone] 1) in nearly all species, in agreement with previous reports (Kishida et al., 2015).

**Figure 3:**
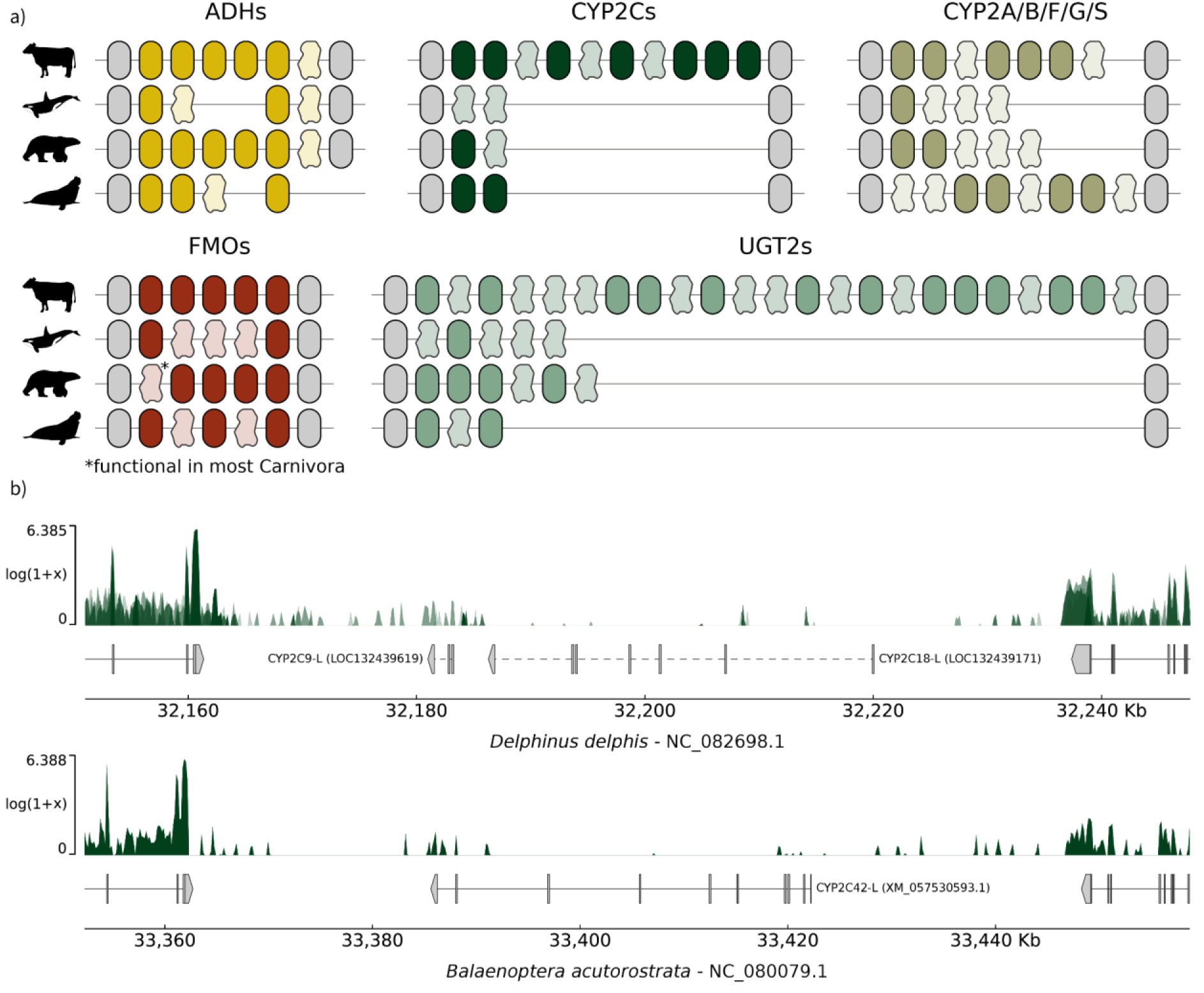
Overview of some of the most dramatically affected biotransformation gene clusters. **a)** Synteny maps of the ADH, CYP2C, CYP2A/B/F/G/S, FMO and UGT2 gene clusters. Each synteny is bordered, in gray, by the conserved genes defining each cluster. The remaining colored boxes and irregular shapes represent coding genes and pseudogenes of each subfamily, respectively. **b)** Liver gene expression within the CYP2C locus in *Delphinus delphis* and *Balaenoptera acutorostrata*. For each species, the top track graphs the log1p value of raw reads per bin, and the bottom track the longest transcript of each gene, including chromosomic position and accession numbers for the genomic region. In each side, a fragment of the neighbouring genes is shown, enclosing the CYP2C pseudogenes, labelled with the protein product and accession number. *Delphinus delphis*’ coverage overlays the result for all four used RNA-sequencing runs. The CYP2C annotations for *Delphinus delphis*, dashed, are set as pseudogenes in the assembly; the annotation for *Balaenoptera musculus*’ is set as coding, albeit with a low-quality protein product.

The oxygenases are one of the most variable groups of chemical defensome genes, with many different gene families and subfamilies and a relatively high variance in the number of functional genes per (sub)family (Fig. 2). While CYPs are often most associated with oxygenase biotransformation activity, we also investigated alcohol and aldehyde dehydrogenases (ADHs and ALDHs), FMOs, and prostaglandin-endoperoxide synthases (PTGSs). ALDHs and PTGSs are the only gene families where we did not observe differences in the number of functional genes between lineages. When looking at the ADH gene family, only ADH1 and ADH5 could be reliably identified in all investigated lineages (Fig. 3a & S2). In cetaceans, these two are generally the only functional ADHs that are found in the principal ADH gene cluster.. ADH4, ADH7, and sometimes ADH6 pseudogenes are detected, and these show generally low expression in cetacean liver tissue (Fig. S3). Other cetartiodactyls generally have ADH1, ADH4, ADH5, and one or multiple ADH6s. Carnivorans have the same genes, but generally have only one functional ADH6 gene. In pinnipeds, however, ADH6 shows evidence of pseudogenization in five out of the nine investigated species. Afrothers often have multiple ADH1 copies, ADH4, and ADH6, although ADH6 is lost in elephants and the Florida manatee. Outside of the main ADH cluster, we detected a second putatively functional ADH5-like gene in eight out of 21 cetacean genomes. In addition, dihydrodiol dehydrogenase (DHDH) was detected in all lineages except cetaceans, where the gene is either missing or pseudogenized. In the FMOs, we observed a noticeable difference between marine and terrestrial lineages (Fig. 3a & S4). All main FMOs found in the human genome (FMO1-5), and FMO6 (which is pseudogenized in humans) are found in all lineages, but nearly all lineages show pseudogenization of one or more FMOs. In terrestrial cetartiodactyls, all FMOs appear to be functional. In contrast, cetaceans are missing most FMOs, and only FMO3 is reliably labelled as functional. Alignment of the human FMO4 to Cetartiodactyla FMO4 *loci* revealed frameshift mutations and premature stop codons in both cetaceans and some terrestrial cetartiodactyls. However, these mutations are confined to the end of the final exon, and are unlikely to affect the function of the gene. This is also supported by the RNA expression patterns, which show similar levels of expression in FMO3 and FMO4 in liver tissue of multiple cetacean species (Fig. S5). FMO1 on the other hand, shows more evident signs of pseudogenization in cetaceans, with several inactivating mutations and low gene expression seen for most species. For FMO2, disruptive mutations were only consistently observed in toothed whales, while baleen whales showed at most one. Nevertheless, FMO2 expression is very low in all investigated baleen whales, but as this is predominantly a pulmonary FMO this was to be expected (Dolphin et al., 1998). FMO6 is generally poorly conserved where it is either not detected or only a short fragment is recovered. FMO5 is an outlier, as it is located separately on the genome, and not in the same gene cluster as FMO1-4. In all cetaceans and a handful of terrestrial cetartiodactyls the FMO5 gene is duplicated, while most other species only have one copy. However, an additional FMO5-like gene is detected in many, but not all, terrestrial cetartiodactyls, carnivorans, and afrothers. This FMO5-like gene is located in a known second cluster of FMO genes. The genes in this cluster are either pseudogenized or lost in humans, but they are still functional in mice (Hernandez et al., 2004). The six FMOs are generally well conserved in terrestrial carnivorans and afrotheres as well, while pseudogenization of FMO1 and FMO6 was detected in pinnipeds, and of FMO3, FMO5, and FMO6 in the manatee.

The CYPs are the largest group of phase I oxygenases, and show some of the largest differences in functional gene counts between lineages. The CYP1 family is the only where we did not observe any differences between lineages, as all of them contain three functional CYP1 genes: CYP1A1, CYP1A2, and CYP1B1. In the CYP2 family, the largest of the investigated CYP families, we observed a significant reduction in the number of functional genes in marine mammals compared to their close relatives (Fig. 2). Cetaceans have the fewest CYP2s: only between seven and eight functional genes compared to nearly 25 CYP2s on average in other cetartiodactyla. Similarly, pinnipeds (12-13 on average) and the manatee (14) have fewer CYP2s than other carnivorans (16 on average) or afrotheres (24-25). Loss of CYP2s in marine mammals is found across different CYP2 subfamilies, with a mixture of shared and lineage-specific losses (Table 1). The CYP2A and CYP2D families are reduced in all three investigated marine mammal lineages. Most cetartiodactyls have one CYP2A gene (with exception of the hippopotamus, which has 3), while cetaceans have no functional CYP2As. Terrestrial carnivorans and afrotheres have on average around two CYP2As, while only one functional copy is found in pinnipeds, and none in the manatee. Similarly, cetaceans and pinnipeds generally have no functional CYP2D genes, although CYP2D does not show signs of pseudogenization in the baleen whales. Their terrestrial counterparts generally have one (carnivorans) or two (cetartiodactyls) CYP2Ds. The manatee has retained a functional copy of CYP2D, but this is less than the two or more copies found in other afrotheres. While CYP2Ds have their own genomic locus, CYP2As are part of a larger cluster of CYP2 genes of different CYP2 subfamilies (Fig. 3a & S6). In humans, this cluster consists of three CYP2A genes, and one CYP2B, CYP2S, and CYP2F gene. In addition, the cluster contains multiple known pseudogenes of CYP2As, CYP2Bs, CYP2Gs, CYP2Fs, and CYP2Ts (Fig. S6). In cetaceans, this gene cluster is heavily reduced, and only CYP2S is present and functional in most genomes (Fig. S7). A CYP2G gene is found in all baleen whales and three toothed whales, but it is unknown whether this gene is functional or not. CYP2As and CYP2Bs are generally missing from cetacean genomes, or are clearly pseudogenized. CYP2Fs are still present in most cetacean genomes, but are often pseudogenized and show low expression in liver tissue. In other marine mammals, this gene cluster is less reduced, but still contains generally fewer functional genes than in their terrestrial relatives (Table 1). For example, pinnipeds have lost CYP2S and have fewer functional CYP2Bs, while the manatee has no functional CYP2Fs.

**Table 1:** Average number of CYP2 subfamilies members in different mammal lineages. The first five rows (CYP2A-CYP2S) represent CYP2 subfamilies that are present in the same gene cluster. *: CYP2D is retained specifically in the Mysticeti lineage. **: The CYP2Fs that are not predicted as pseudogenes are divergent from most other CYP2Fs, making pseudogene assessment difficult.

|  | Cetacea | Other<br>Cetartiodactyla | Pinnipedia | Other<br>Carnivora | Sirenia | Other<br>Afrothera |
| --- | --- | --- | --- | --- | --- | --- |
| **CYP2A** | 0 | 1.04 | 1.11 | 2.03 | 0 | 1.83 |
| **CYP2B** | 0 | 2.54 | 0.89 | 1.68 | 2 | 1.5 |
| **CYP2F** | 0.57** | 1 | 2 | 1.91 | 0 | 1.67 |
| **CYP2G** | 0.05 | 0.8 | 1 | 0.82 | 1 | 0.75 |
| **CYP2S** | 0.86 | 0.96 | 0 | 0.88 | 1 | 1 |
| **CYP2C** | 0.95 | 7.08 | 2.56 | 2.21 | 5 | 5.5 |
| **CYP2D** | 0.24* | 2 | 0 | 0.94 | 1 | 2.33 |
| **CYP2E** | 0.95 | 1 | 1.22 | 1.06 | 0 | 0.67 |
| **CYP2J** | 1 | 4.38 | 0.78 | 1 | 0 | 3.83 |
| **CYP2R** | 1 | 0.96 | 1 | 1.06 | 1 | 1 |
| **CYP2U** | 1 | 0.92 | 1.11 | 0.97 | 1 | 1 |
| **CYP2W** | 0.24 | 0.88 | 1 | 1 | 1 | 1 |

Another major CYP2 gene cluster is the CYP2C cluster. This cluster contains the four CYP2C genes in humans (CYP2C8, CYP2C9, CYP2C18, and CYP2C19). This gene cluster is heavily expanded in terrestrial cetartiodactyls, especially in ruminants (Fig. 3a & S8). In cetaceans, this cluster often only contains one or two CYP2C genes, which are nearly always pseudogenized and show low RNA-expression in liver tissue (Fig. 3b & S9). However, cetaceans still have one functional CYP2C gene elsewhere in the genome, a CYP2C23-like gene found in most other cetartiodactyls as well. Both terrestrial and marine carnivorans generally have one or two functional CYP2C genes in this cluster, and a similar CYP2C23-like gene elsewhere in the genome (in tandem with CYP2E). Lastly, afrotheres (including the manatee) have 3-5 CYP2Cs in the cluster, and some have an additional CYP2C23-like gene in the same location as the cetartiodactyls. Other CYP2Cs that are found outside the two main CYP2 clusters are CYP2E, CYP2J, CYP2R, CYP2U, and CYP2W (Table 1). CYP2E is generally well conserved in all species, although it contains a stop codon very early on in the first exon of the two genes in the manatee. CYP2J is expanded in terrestrial cetartiodactyls and afrotheres, but lost in the manatee. The other lineages have mostly one functional copy. CYP2R, CYP2U, and CYP2W are conserved in nearly all investigated species, although CYP2W is pseudogenized in the majority of cetaceans.

The number of CYP3s also varies between and within the investigated lineages. Nearly all investigated cetaceans have two CYP3 genes, while other artiodactyls tend to have four. In the carnivorans, we see a big distinction between feliforms, who have only two CYP3s, and caniforms, who generally have four CYP3s. The number of CYP3s in pinnipeds is difficult to assess due to generally poor genome assembly quality at this locus. We could detect four CYP3s in the California sea lion (*Zalophus californianus*), but all other investigated pinnipeds show fewer CYP3s (generally only one or two complete copies). The CYP3 locus in afrothers is in a different genomic location, and contains between two and four CYP3s, depending on the species. The number of functional CYP4s is lower in all three marine mammals compared to their terrestrial relatives (Fig. 2). The lower number of CYP4s is due to fewer functional CYP4As and loss of CYP4B in all three marine lineages. CYP4X is also lost in cetaceans and the manatee, but is pseudogenized in only three out of nine investigated pinnipeds. Cetaceans also have significantly fewer CYP4Fs, but the difference in other marine lineages is less pronounced. Lastly, CYP4V is retained in all investigated lineages, except in the manatee.

### Phase II biotransformation

In the second phase of biotransformation, hydrophilic groups are transferred from endogenous compounds to the xenobiotics to form more water-soluble and inactive compounds that can be excreted by the body. The main type of enzymes involved in this process are transferases. They encompass some large gene families where we observed substantial variation between and within lineages (Fig. 2). In terms of *N*-acetyltransferases, most lineages have on average eight functional genes. The aralkylamine *N*-acetyltransferase (AANAT) is, however, duplicated in most terrestrial cetartiodactyls, pseudogenized in nearly all whales (nearly exclusively in toothed whales), and not detected in the manatee. In afrotheres (including the manatee), NAT1 is duplicated at least once, and even multiple times in some species. NAT8 is generally poorly conserved, with pseudogenes found in many cetartiodactyls (including all cetaceans), many pinnipeds, and the manatee. NAT6 is missing in some cetaceans, and shows possible pseudogenization in the species where it is present and in the manatee.

The gene inventory of the three largest transferase gene families (GST, SULT, and UGT) show a somewhat similar pattern in terms of distribution: large variations in the number of genes within lineages, and cetaceans (and to a lesser extent other marine lineages) who have generally fewer functional genes (Fig. 2). Functional GSTs are present in fewer numbers in cetaceans (ten genes on average) than in other lineages (18 genes in other cetartiodactyls, 15 in carnivorans, and 17 in afrotheres). Interestingly, the average number of functional GSTs is higher in caniforms (+-16) than in feliforms (+- 12). The lower number of GSTs in cetaceans is due to a reduction in the GSTA, GSTM, and GSTT subfamilies (Fig. 2). The three microsomal GSTs (MGST1-3) are generally well-conserved across the investigated species, although the manatee has lost MGST1, and a large proportion of cetaceans have lost MGST2. Similarly, we observed a reduced number of sulfotransferases (SULTs) in cetaceans, but also in other marine mammals. Terrestrial cetartiodactyls and carnivorans have on average 9-10 SULTs, while cetaceans and pinnipeds only have 6-7. The manatee has slightly more (eight), but this is still lower than the average for other afrotheres (eleven). The observed lower numbers of SULTs are almost exclusively due to a reduction in the SULT1C subfamily and loss of SULT6B in all marine mammals, loss of SULT2A in cetaceans, and loss of SULT1E1 in pinnipeds (Fig. 2). Lastly, the UGT family is the largest, but also the most diverse phase II biotransformation gene family. The number of detected functional UGTs in terrestrial cetartiodactyla ranges from eight to 33 genes, with an average of nearly 18 genes. In contrast, cetaceans have on average only three to four UGTs, with at most six genes (Fig. 3a). The majority of UGTs are UGT2s which are located in the same gene cluster (Fig. S10). Similar to the CYP2C gene cluster, this cluster is heavily expanded in terrestrial cetartiodactyls (and especially ruminants), and significantly reduced in cetaceans. While terrestrial cetartiodactyls have on average ten UGT2s in this cluster, we could rarely detect more than one functional copy in cetaceans, which shows signs of being expressed in liver tissue (Fig. S11). Pinnipeds have two functional genes in the cluster, while generally more are found in terrestrial carnivorans. In the manatee, all three identified genes show signs of pseudogenization, while the gene cluster have two to three functional genes in other afrotheres (except both included elephant species, which each have over ten genes in the cluster). Similarly, UGT1s have expanded in terrestrial cetaceans and afrotheres, while only copy is present in cetaceans and two in carnivorans. UGT3s are lost in cetaceans and the manatee, while UGT8 is present in all investigated lineages.

### Transporters

Membrane transporters of the chemical defensome can be broadly separated into two groups: the ABC-transporters, and the ion-metal transporters (solute carriers or SLCs). Most ABC and SLC gene families have similar numbers of functional genes in the different lineages, although terrestrial cetartiodactyls have elevated numbers of functional genes in the two largest gene families: ABCC and SLC22 (Fig. 2). The higher observed number of ABCCs in terrestrial cetartiodactyls is mainly due to an expansion of ABCC4 and ABCC4-like genes. While nearly all investigated species have only one ABCC4 gene, ruminants have on average more than six ABCC4 or ABCC4-like genes. A large part of this gene expansion is local duplication of the original ABCC4 gene, but additional ABCC4-like genes elsewhere in the genome also contribute to the increased number of total ABCCs in the ruminants. In addition to having fewer ABCC4s, cetaceans also have lost both ABCC11 and ABCC12. Interestingly, the pygmy sperm whale (*Kogia breviceps*) has 20 ABCC6-like genes in addition to the ABCCs found in other cetaceans. However, these copies include only exons 7-11 out of a total of 31 exons for ABCC6 (not shown), coding for proteins of only 250 to 260 amino acids, while most ABC transporters are around 1500 amino acids. This indicates these genes might not be functional. Outside of the ABCCs, only few differences between lineages were observed. Cetaceans and the manatee have a pseudogenized ABCB5, and pseudogenization of ABCG5 and ABCG8 is common in pinnipeds and afrotheres. These two latter genes are found adjacent in the genomes and are nearly exclusively lost as a pair. Indeed, the products of these two genes are known to function as heterodimers (Graf et al., 2003).

Similar to the ABC transporters, most solute carrier (SLC) families have similar numbers of functional genes in each lineage, except for the highly diverse SLC22 family and, to a lesser extent, the SLC47 family (Fig. 2). Most species have around 20 SLC22s, but the average number is slightly lower in cetaceans (16.7), and slightly higher in terrestrial cetartiodactyls (22.5). These deviations are mostly because of loss of SLC22A11, SLC22A12, SLC22A13, and SLC22A31 in cetaceans, and expansion of SLC22A9 and SLC22A10 in other cetartiodactyls. SLC47A1 is also absent or pseudogenized in some cetaceans. In terms of SLCOs, gene loss is only observed in the manatee. In this species, SLCO1A2, SLCO1B3, SLCO1C1, and SLCO5A1 are all lost.

### Stress responses (antioxidants, metal response, heat response)

Many stress response genes are also involved in the response to foreign chemicals. Metal-responsive genes are activated when cells are exposed to metals, while heat-shock proteins and antioxidants protect against cellular stress, which might occur during the detoxification process. The antioxidant gene inventory is relatively stable among lineages, apart from some minor differences in the peroxiredoxins (PRDXs) and thioredoxins (TXNs) (Fig. S2). PRDXs tend to be present in more copies in cetaceans and pinnipeds compared to their close relatives, although the variation within each lineage is quite large. In contrast, the manatee has the lowest number of PRDXs of the investigated afrotheres.The higher number of genes in these first two groups is due to duplications of PRDX1 (both lineages), PRDX3 (cetaceans), and PRDX5 (pinnipeds). In addition, cetaceans tend to have more TXNs compared to other cetartiodactyls (Fig. S1). Heat-responsive genes also showed little differences between lineages. The bulk of heat-responsive genes are members of the DNAJ and HSP families, which are present in relatively large numbers (39-58 DNAJs, 23-56 HSPs). DNAJs are very diverse, leading to clustering in many different orthologous groups, often only containing one or two genes per species. Cetaceans and the manatee seem to have slightly fewer DNAJ genes than their respective close relatives, but the averages are relatively close (ranging from 40 to 49 genes on average). The heat-shock proteins (HSPs) are similarly diverse but show no large differences between lineages except a slightly higher average number of HSPs in pinnipeds compared to other carnivorans (36 vs 31). Similar to antioxidants and heat-responsive genes, we observed no large differences in the number of metal-responsive genes between the investigated lineages (Fig. S1). The number of ceruloplasmins (CPs) does show slight variations, where carnivorans and many terrestrial cetartiodactyls have two copies, while cetaceans and afrotheres only have one. The number of ferritins (both heavy chains, FTH, and light chains, FTL) is very variable within lineages, ranging from only a couple to over 20. Lastly, the number of detected functional metallothioneins (MTs) is highly variable within cetaceans, ranging from two to 35 copies.

## Discussion

### A framework for assessing the chemical defensome

In this work we identified the chemical defensome in a total of 97 species in three mammalian lineages: Cetartiodactyla, Carnivora, and Afrotheria. We make use of the increased availability of high-quality reference genomes thanks to large projects such as the Earth Biogenome Project (Lewin et al., 2018) and the Darwin Tree of Life (The Darwin Tree of Life Project Consortium, 2022) to perform a large-scale genomic comparison of the chemical defensome. Using well-established genomic tools and previous knowledge of the chemical defensome, we identified on average 474 genes in the chemical defensome of the investigated species. The number of identified genes is similar to those found using similar methods in fish (446-510 genes; Eide et al., 2021). The genes in the chemical defensome are similarly distributed among the different categories, with the average numbers of phase I oxygenases and reductases, phase II biotransformation enzymes, and transporters falling within or close to the range observed in fish (Table 2). If we compare some of the more prominent gene (super)families (CYPs, AKRs, GSTs, UGTs, SULTs, and ABCs), some invertebrates like copepods show a chemical defensome very similar to that of vertebrates (Roncalli et al., 2025). However, the purple sea urchin (*Strongylocentrotus purpuratus*) and the starlet sea anemone (*Nematostella vectensis*), two of the first species wherein the chemical defensome was described, show a highly elevated number of CYPs (mainly CYP2s), SULTs, and ABCs compared to vertebrates (Goldstone, 2008; Goldstone et al., 2006). The number of GSTs and UGTs is also significantly higher in the purple sea urchin, but not in the sea anemone, highlighting the high diversity of the chemical defensome in invertebrates.

**Table 2:** Overview of the defensome in different lineages and species. For lineages investigated in this study. the average number (rounded to the nearest integer) is given. Sources: (1) this study, (2) Goldstone et al., 2006, (3) Goldstone et al., 2008, (4) Eide et al., 2021, (5) Roncalli et al., 2025, (6) De Marco et al., 2017.

|  | Artiodactyla<br>(1) | Cetacea<br>(1) | Carnivora<br>(1) | Pinnipedia<br>(1) | Afrotheria<br>(1) | Sirenia<br>(1) | Human<br>(*Homo sapiens*)<br>(2,3) | Purple sea urchin<br>(*Strongylocentrotus purpuratus*) (2,3) | Starlet sea anemone<br>(*Nematostella vectensis*)<br>(2,3) | Teleosts<br>(min - max) (4) | Copepods<br>(min - max) (5) | *Anopheles stephensi*<br>(6) |
| --- | --- | --- | --- | --- | --- | --- | --- | --- | --- | --- | --- | --- |
| **Total** | **515** | **446** | **458** | **471** | **487** | **447** | **218** | **423** | **226** | **446-510** |  |  |
| **Oxygenases** | **120** | **67** | **92** | **77** | **117** | **78** |  |  |  | **60-81** |  |  |
| ADH | 6 | 3 | 5 | 5 | 5 | 5 |  |  |  |  |  |  |
| ALDH | 18 | 18 | 19 | 20 | 19 | 16 | 19 | 20 | 21 | 19-22 | 4-9 |  |
| CYP (all) | 44 | 20 | 30 | 23 | 42 | 26 | 57 | 120 | 82 | 29-54 | 22-36 | 44 |
| CYP1 | 3 | 4 | 3 | 3 | 4 | 3 | 3 | 11 |  | 3-5 |  |  |
| CYP2 | 25 | 8 | 16 | 12 | 25 | 14 | 21 | 85 | 39 | 18-40 |  |  |
| CYP3 | 4 | 2 | 3 | 2 | 3 | 4 | 5 | 13 | 20 | 4-4 |  |  |
| CYP4 | 12 | 6 | 8 | 6 | 10 | 5 | 12 | 10 | 3 | 4-5 |  |  |
| FMO | 6 | 4 | 6 | 4 | 7 | 3 | 6 | 16 | 6 | 1-4 | 1-2 |  |
| PTGS | 2 | 2 | 2 | 2 | 2 | 2 |  |  |  |  |  |  |
| **Reductases** | **41** | **30** | **31** | **30** | **35** | **32** |  |  |  | **28-37** |  |  |
| AKR | 17 | 7 | 7 | 6 | 11 | 8 | 8 | 10 | 12 |  | 2-4 | 1 |
| EPHX | 4 | 4 | 4 | 4 | 4 | 3 | 2 | 5 | 1 |  |  |  |
| HSD | 16 | 16 | 16 | 16 | 15 | 16 |  |  |  |  |  |  |
| HSDL | 2 | 2 | 2 | 2 | 2 | 3 |  |  |  |  |  |  |
| NQO | 2 | 1 | 2 | 2 | 3 | 2 | 2 | 0 | 0 | 1-10 |  |  |
| **Transferases** | **57** | **27** | **46** | **41** | **57** | **37** |  |  |  | **39-76** |  |  |
| AANAT | 2 | 0 | 1 | 1 | 1 | 0 |  |  |  |  |  |  |
| GST | 18 | 10 | 15 | 15 | 17 | 15 | 21 | 38 | 18 | 9-19 | 16-28 | 22 |
| MGST | 3 | 2 | 3 | 3 | 4 | 2 | 3 | 12 | 5 |  |  |  |
| NAT | 6 | 5 | 7 | 7 | 9 | 6 | 2 | 1 | 0 |  |  |  |
| SULT | 10 | 6 | 10 | 7 | 11 | 8 | 13 | 73 | 22 |  | 3-8 |  |
| UGT | 18 | 4 | 10 | 8 | 15 | 6 | 13 | 50 | 9 | 11-31 | 5-11 | 13 |
| **Transporters** | **95** | **79** | **83** | **83** | **84** | **74** |  |  |  |  |  |  |
| ABC (all) | 36 | 24 | 24 | 24 | 24 | 22 | 48 | 65 | 65 | 27-32 | 20-37 |  |
| ABCB | 9 | 8 | 8 | 9 | 9 | 8 | 11 | 12 | 7 | 8-10 |  |  |
| ABCC | 22 | 11 | 11 | 11 | 11 | 11 | 12 | 30 | 36 | 12-14 |  |  |
| ABCG | 5 | 5 | 5 | 4 | 4 | 3 | 5 | 9 | 6 | 7-8 |  |  |
| SLC (all) | 59 | 55 | 59 | 59 | 60 | 52 |  |  |  | 42-59 |  |  |
| SLC22 | 23 | 17 | 20 | 20 | 22 | 20 | 11 | 30 | 17 |  |  |  |
| SLC30 | 9 | 10 | 10 | 10 | 10 | 10 |  |  |  |  |  |  |
| SLC31 | 2 | 2 | 2 | 2 | 2 | 2 |  |  |  |  |  |  |
| SLC39 | 14 | 14 | 14 | 14 | 14 | 14 |  |  |  |  |  |  |
| SLC47 | 2 | 2 | 3 | 3 | 2 | 0 |  |  |  |  |  |  |
| SLCO<br>(OATP) | 9 | 10 | 10 | 10 | 10 | 6 |  | 30 |  |  |  |  |

A significant part of the analysis pipeline used in this work involves pseudogene prediction. Identifying genes that might no longer be functional is an important step, as it allows the detection of ongoing gene loss, even if remnants of the gene still remain. Examining only the presence and absence of genes may overlook a significant part of lost genes, as is demonstrated by PXR and CAR, which are still present in the genome but show clear signs of erosion. The high number of identified putative pseudogenes in the chemical defensome (up to 18%) highlights the importance of considering pseudogenes when comparing gene inventories between species. However, pseudogene prediction is far from straightforward, and the results need to be assessed critically. One important factor is the number and location of inactivating mutations (e.g. premature stop codons or frameshifts) to consider a pseudogene (Alves et al., 2020). In our work, we used relatively permissive restraints, requiring only one inactivating mutation in the gene’s coding sequence. While this might erroneously label genes as pseudogenes (for example in the case of FMO4), we manually inspected these alignments, and in some cases used RNA-seq expression data, to make a call. This led us to identify a specific case of wrongly labelled pseudogenes, namely genes that contain selenocysteins. These special amino acids are coded by TGA codons which are normally used as stop codons. In these cases, the gene would be labelled as a pseudogene even though it is likely functional (e.g. this was the case in GPX3, 4, and 6; (Brigelius-Flohé and Maiorino, 2013)). Pseudogene prediction also requires a suitable reference protein for performing the alignment. In cases where no reference protein is available, we cannot assess whether a gene is functional based on the sequence alone. Additionally, in cases of one-to-many orthology, aligning a single reference gene to all orthologs might incorrectly classify genes as pseudogenes based on low sequence identity or coverage, even though these genes may simply have diverged following duplication. This is especially true within the scope of our analyses, as a considerable number of defensome gene clusters underwent tandem duplication and pseudogenization. Nevertheless, the majority of gene losses observed in the chemical defensome correspond to genes that are either absent from the genome or exhibit clear evidence of pseudogenization.

### Significant reduction of the chemical defensome in cetaceans

While we observed gene loss in all three investigated marine mammal lineages, the reduction in the number of chemical defensome-related genes in the cetacean lineage is the most striking. These gene losses were observed in nearly all phases of chemical detoxification from nuclear receptors, over phase I and phase II biotransformation, to transporters. The losses in transcription factors are the fewest, but possibly the most impactful. We confirmed the loss of both PXR (NR1I2) and CAR (NR1I3) in cetaceans, as previously reported in multiple studies (Hecker et al., 2019; Wagner et al., 2022). Both proteins are key xenobiotic receptors which recognize a broad range of ligands, and regulate a set of overlapping genes involved in phase I and phase II of drug metabolism, as well as transporters (Tolson and Wang, 2010). When looking specifically at genes that are known to be regulated by PXR or CAR, we notice many of them are indeed lost in cetaceans (Table 3). Many of the CYP families regulated by PXR and/or CAR are either lost, or contain significantly fewer copies in cetaceans than in other cetartiodactyls. Similarly, transferases regulated by PXR/CAR in humans are also present in fewer copies in cetaceans. For some SULTs, evidence points towards regulation by PXR/CAR in mice and rats, but only SULT2A is lost in cetaceans (Tolson and Wang, 2010). In contrast to phase I and II biotransformation enzymes, transporters regulated by PXR/CAR seem to be less affected in cetaceans. Only in the ABCC4-family we see significantly fewer genes in cetaceans than in other cetartiodactyla, but this is more likely due to expansion of this gene family in ruminants. A similar case can be made for the AKR1C-family, where terrestrial cetartiodactyls might have expanded their gene repertoire compared to other lineages.

**Table 3:** Known target genes of NR1I2 (PXR) and NR1I3 (CAR) and their presence in marine mammals. PXR and CAR targets are derived from Tolsen and Wang, 2010. Lost genes mean that no functional copies of the gene or gene families are found in the lineage, reduced means that that lineage contains fewer functional genes in that gene family than the terrestrial relative lineage.

| Gene (family) | PXR target? | CAR target? | Gene loss |  |  | Note |
| --- | --- | --- | --- | --- | --- | --- |
|  |  |  | Cetacea | Pinnipedia | Sirenia |  |
| CYP1A1 | - | Mouse |  |  |  | Retained in all marine mammals |
| CYP1A2 | - | Mouse |  |  |  | Retained in all marine mammals |
| CYP2A | - | Mouse (CYP2A4) | X | X | X | Lost in Cetacea, reduced in other marine mammals |
| CYP2B | Human (CYP2B6); Rat (CYP2B1/2); Mouse (CYP2B10) | Human (CYP2B6); Rat (CYP2B1/2); Mouse (CYP2B10) | X | X |  | Lost in Cetacea and most Pinnipedia |
| CYP2C | Human (CYP2C8, CYP2C9, CYP2C19) | Human (CYP2C8, CYP2C9, CYP2C19); Mouse (CYP2C29, CYP2C37) | X |  |  | Reduced in Cetacea |
| CYP3A | Human (CYP3A4, CYP3A7); Mouse (CYP3A11); Rat (CYP3A2, CYP3A23) | Human (CYP3A4); Mouse (CYP3A11) | X | X |  | Reduced in Cetacea and Pinnipedia |
| CYP4F12 | Human | - | X |  |  | Lost in Cetacea |
| AKR1C | Human (AKR1C1, AKR1C2) | - | X |  | X | Reduced in Cetacea and Sirenia |
| AKR1B | Mouse (AKR1B7) | Mouse (AKR1B7) | X |  | X | Reduced in Cetacea and Sirenia |
| ALDH1A1 | Mouse | Mouse |  |  |  | Retained in all marine mammals |
| UGT1A | Human (UGT1A1, UGT1A3, UGT1A6); Mouse (UGT1A1, UGT1A9) | Human (UGT1A1); Mouse (UGT1A1, UGT1A6, UGT1A9) | X |  |  | Reduced in Cetacea |
| UGT2B | Mouse (UGT2B5) | Mouse (UGT2B1) | X | X | X | Reduced in all marine mammals |
| GSTA | Human (GSTA1); Mouse (GSTA1/2) | Mouse (GSTA1/2) | X |  | X | Reduced in Cetacea and Sirenia |
| SULT1A1 | Rat | - |  |  |  | Retained in all marine mammals |
| SULT1B1 | Rat | - |  |  |  | Retained in all marine mammals |
| SULT1E1 | Mouse | Mouse |  | X |  | Lost in Pinnipedia |
| SULT2A | Mouse (SULT2A1, SULT2A2); Rat (SULT2A3/4) | Mouse (SULT2A1, SULT2A2) | X | NA |  | Lost in Cetacea; Not present in Carnivora |
| ABCB1 | Human, Mouse | Human, Mouse |  |  |  | Retained in all marine mammals |
| ABCC1 | - | Mouse |  |  |  | Retained in all marine mammals |
| ABCC2 | Human, Mouse | Human, Mouse |  |  |  | Retained in all marine mammals |
| ABCC3 | Mouse | Human, Mouse |  | X |  | Lost in some Pinnipedia |
| ABCC4 | - | Human, Mouse | X |  |  | Reduced in Cetacea |
| SLCOA1 | Mouse, Rat | - |  |  | X | Lost in Sirenia |

To further investigate the hypothesis that some of the observed losses in downstream genes might be related to the loss of PXR and CAR, we investigated the functional and physical connectivity between genes of the chemical defensome (Fig. 4). The chemical defensome is highly functionally connected, with 313/406 genes connected in a single network. Clustering based on confidence of the connections between genes revealed different clusters related to different stages and functions of the chemical defensome, with NR1I2 and NR1I3 sitting as hubs between different clusters. Cetacean gene loss is most prevalent in the tightly connected clusters of phase I and phase II biotransformation enzymes, especially genes that are regulated by NR1I2 and/or NRI1I3, and in xenobiotic and organic anion transporters. Cetacean gene loss is less common in other clusters, further supporting the connection between the loss of NR1I2 and NR1I3 and the loss of biotransformation genes. Cetacean gene loss is also found in just over 30% of genes outside the main network (29/93 genes). The majority of these genes are also phase I and phase II biotransformation genes or transporters, showing a broad impact on the chemical defensome as a whole.

**Figure 4:**
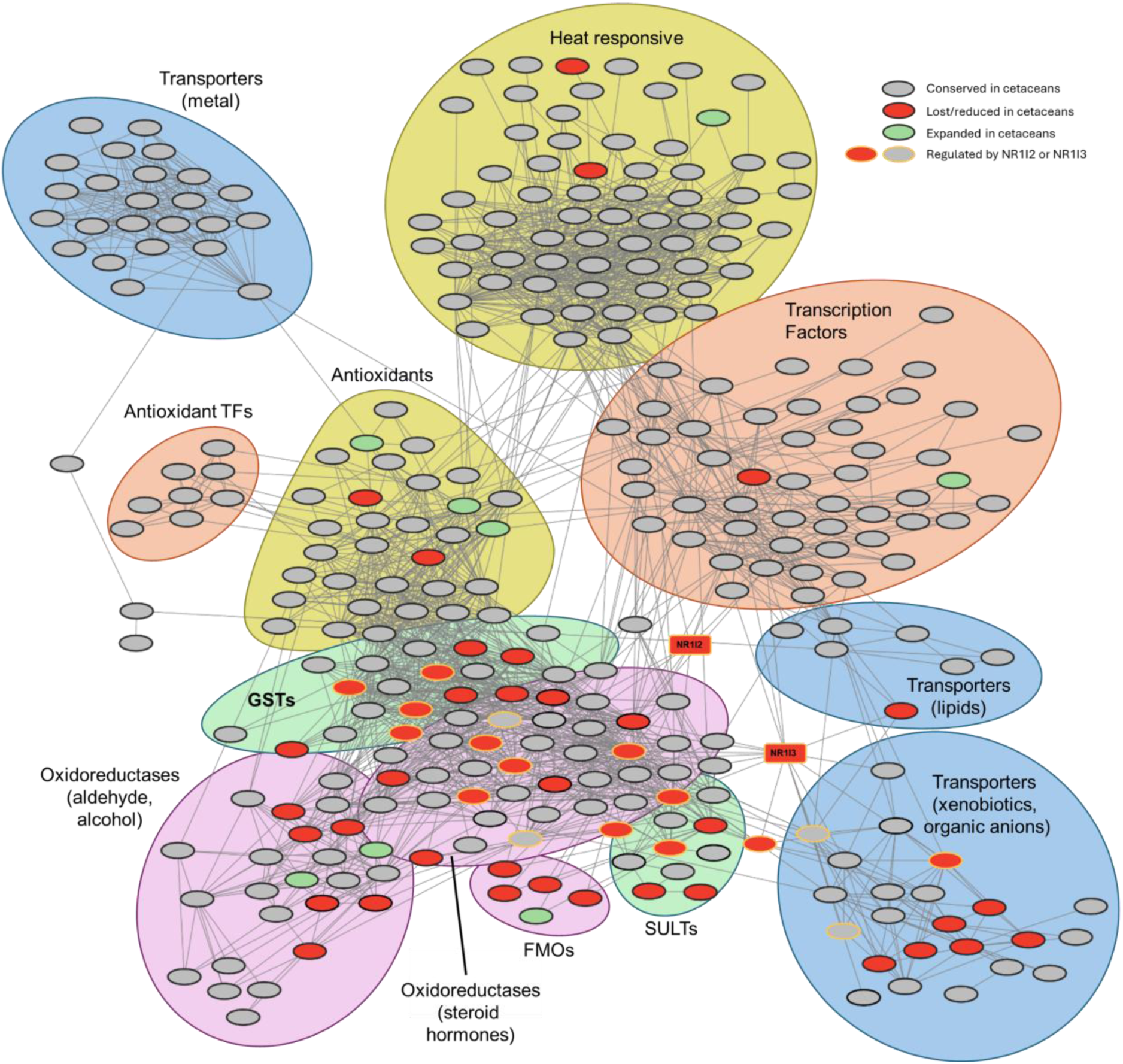
STRING network of the chemical defensome. Nodes represent genes of the chemical defensome with an equivalent in the human genome in the STRING database (https://string-db.org/; Szklarczyk et al., 2022), and are connected in the main network (313 genes). Singleton genes (genes without connections; 86 genes) and genes in small networks (<= 3 connected genes; 7 genes) are not shown. Edges represent functional interactions/correlations between proteins, based on experimental data, database information, co-expression data, gene neighbourhood data, gene fusion data, co-occurrence data, and textmining.Only edges with high confidence (>= 0.7) in the STRING database are shown. The network was clustered using Edge-Weighted Spring Embedded Layout followed by overlap reduction, implemented in Cytoscape (v. 3.10.4; Shannon et al., 2003), using the STRING edge confidence as weights. Nodes colors correspond to conservation status in cetaceans, compared to other cetartiodactyla: red - lost or reduced in cetaceans, green - expanded in cetaceans, or lost in other cetartiodactyla; grey - no difference between cetaceans and other cetartiodactyla. Nodes with a yellow colored border indicate genes that are known to be regulated by PXR (NR1I2) or CAR (NR1I3) in humans. PXR and CAR are indicated as red squares in the network. Subnetworks/clusters in the network are highlighted according to the functions of the containing genes, based on manual inspection of the gene and functional enrichment analysis in STRING.

Since both nuclear receptors are known to share many target genes (Tolson and Wang, 2010), it is unclear whether loss of one or both of these transcription factors is correlated with the observed large scale losses in the chemical defensome of cetaceans. While cetaceans are the only mammals to have lost both PXR and CAR, loss in one of both transcription factors has been reported in multiple other mammalian lineages. In bats (Chiroptera), loss of CAR is observed in the Yangochiroptera suborder, and loss of both PXR and CAR is also documented in the Eulipotyphla order of insectivores (Hecker et al., 2019). Preliminary investigation of the chemical defensome in these lineages using the same methods we applied for the marine mammals showed no large-scale losses as observed in cetaceans (data not shown). Chiropterans in general did not show reduced numbers of oxygenases, reductases, transferases, or transporters compared to carnivorans, and neither did we observe a reduced chemical defensome in the suborders Yangochiroptera (who have mostly lost CAR) and Yinpterochiroptera (who have both PXR and CAR). In Eulipotyphla, the situation is more unclear. Loss of PXR is only found in the Soricidae family, while CAR is only lost in the Talpidae. However, neither of these losses seems to be lineage-specific (PXR is lost in 2/3 Soricidae, CAR in 1/3 Talpidae), and the low number of genomes in both lineages makes large-scale comparison of the chemical defensome difficult. Outside of Mammalia, loss of PXR has been documented for several teleost fish species (Eide et al., 2018). Over half of the 76 investigated teleost genomes showed loss of PXR, including almost all members of the Gadiformes order. Since fish diverged from the vertebrate lineage before the duplication of PXR leading to the formation of CAR, they only possess PXR (Handschin et al., 2004). Further investigation into the whole chemical defensome in five teleost model species did not reveal any compensatory mechanisms nor loss of downstream chemical defensome genes in the two species that have lost PXR (Atlantic cod, *Gadus morhua*, and three-spined stickleback, *Gasterosteus aculeatus*) (Eide et al., 2021). However, an increased number of AHR-reponsive regulatory elements were detected upstream of CYP3A in the Atlantic cod, suggesting an extended regulatory role for the aryl hydrocarbon receptor (AHR) in the chemical defensome in fish (Eide et al., 2018).

Within the cetacean lineage, we found some small differences in the chemical defensome between baleen whales (Mysticeti) and toothed whales (Odontoceti). For example, CYP2D6, CYP2G1, AANAT, and FMO2 all show preferential pseudogenization in toothed whales, while we did not observe any genes that are specifically lost in baleen whales. In some cases, it is not certain that the retained gene in the baleen whales is actually still functional. For example, a CYP2G-like gene is only found in three toothed whale genomes, while all investigated baleen whale genomes contain the gene. Alignment to the CYP2G1 sequence of mice shows these genes do not seem to contain any disrupting mutations. However, we did not detect any expression of this gene in cetacean liver tissue, which is in accordance with previous reports of the gene’s function in the olfactory mucosa (Hua et al., 1997). In case of AANAT, reports show a shared inactivating premature stop codon near the end of the first exon in all cetaceans, but different additional mutations between baleen and toothed whales (Lopes-Marques et al., 2019). Further investigation into our pseudogene alignments revealed that baleen whale AANAT was labelled as functional because both the premature stop codon in exon 1 and the altered splice site in exon 3 could be avoided by using an nearby alternative splice site. In contrast, the AANAT gene in toothed whales contains different disrupting mutations which can not easily be avoided. Whether the alternative splice sites in baleen whale AANAT can effectively be used, and thus if the gene is potentially functional in baleen whales or not, is up for debate. *In vitro* minke whale (*Balaenoptera acutorostrata*) AANAT activity was nearly undetectable, although the recombinant protein was based on the predicted CDS and did not account for the possibility of alternative splicing variants (Yin et al., 2021). In any case, both the CYP2G and AANAT examples reveal different rates of gene erosion between toothed and baleen whales.

Despite the large number of gene losses and gene family reductions, we also observed gene family expansions in cetaceans. Most of these gene expansions (or gene retentions) are, however, relatively small. For example, we observed an increase in the number of thioredoxins (TXNs) and peroxiredoxins (PRDXs) in cetaceans, but in both cases cetaceans only have one additional gene on average. Both gene families are antioxidants, a group of genes responsible for protecting the cell from oxidative stress. Gene expansion and molecular changes in antioxidants have been linked to protection against hypoxia in long and deep-diving animals such as cetaceans and pinnipeds (Allen and Vázquez-Medina, 2019; Selleghin-Veiga et al., 2024), and would explain also the expansion of TXNs observed in pinnipeds. Additionally, the higher number of antioxidants could help against the possible negative effects from the observed loss of NQO1 in cetaceans. NQO1 plays an important role in detoxification of reactive oxygen species (ROS) by removing quinones from the cell (Torrente et al., 2020), and the expansion of TXNs and PRDXs in cetaceans could help reduce cellular stress related to increased ROS-levels owing to the loss of NQO1.

### Convergent and independent defensome losses in other marine mammal lineages

Despite having retained both PXR and CAR, we also observed significant gene loss in the other investigated marine mammal lineages (Pinnipedia and Sirenia), particularly in the phase I and phase II biotransformation enzymes such as CYPs, FMOs, and SULTs. Interestingly, most of the gene losses that are shared between multiple marine mammal lineages, are genes or gene families that are not directly regulated by PXR or CAR. For example, FMOs are not known to be regulated by either of these transcription factors while we observed loss of FMOs in all marine mammals (although FMO6 is the only FMO lost in all marine lineages). Loss of SULTs in pinnipeds has been described in the literature before (Kondo et al., 2023), and here we found similar gene loss in cetaceans and sirenians. We found fewer SULT1s in all marine mammals, and nearly all investigated marine mammals have lost SULT6. While the function of SULT6 is still mostly unknown, SULT1s are known to play an important role in detoxification of xeno-and endobiotic compounds (Coughtrie, 2016). Their regulation is still not fully elucidated, with evidence pointing to regulation by multiple nuclear receptors, such as PXR, CAR, PPAR, VDR, FXR, and GR (Alnouti and Klaassen, 2008; Runge-Morris et al., 2013; Tolson and Wang, 2010). However, the SULTs that are specifically lost in marine mammals (SULT1D1, SULT1Cs, and SULT6) do not show direct evidence of regulation by PXR or CAR, and generally are poorly or not at all functionally connected to the rest of the chemical defensome (Fig. 4). Lastly, marine mammals share a general decrease in the number of functional CYP2s and CYP4s in their genomes. Interestingly, reduction in the number of CYP2Cs, a group of CYPs that is often regulated by PXR and/or CAR, is only observed in cetaceans, while CYP2 subfamilies that are lost or reduced in all marine mammal lineages (e.g. CYP2A and CYP2D) are not directly linked to PXR or CAR regulation (although some evidence points towards regulation of CYP2A4 in mice (Tolson and Wang, 2010)). Reduction of CYPs in sirenians is in congruence with previous research (Watanabe et al., 2023), although a similar screening for CYPs in the Carnivora lineage did not observe reduction of CYP2Bs and loss of CYP2D as we report here (Kondo et al., 2022). In both cases, the observed gene loss is due to the presence of inactivating mutations, which might not have been considered in previous studies.

While not part of any of the three main marine mammal lineages, the polar bear (*Ursus maritimus*) and the sea otter (*Enhydra lutris*) could still be considered marine mammals. Both species belong to the Carnivora lineage, and spend a significant amount of their life in a saltwater environment (Würsig et al., 2018). Similar to other carnivorans, they have both retained NR1I2 and NR1I3, but have diverged more recently from their terrestrial ancestors than the pinnipeds (e.g. polar bears diverged from brown bears approximately 1-1.6 Mya) (Agnarsson et al., 2010; Lille-Langøy et al., 2015; Sun et al., 2024). In neither sea otter or polar bear we observed gene loss or gene family reduction on the scale found in other marine mammals. However, the sea otter has the lowest number of phase I and phase II biotransformation genes of all mustelids. This could indicate first steps towards a reduction in the chemical defensome due to their shift to the marine environment, although their chemical defensome is still considerably larger than most pinnipeds. Polar bears have a very similar chemical defensome to other bears, which is not surprising given the very recent divergence of this species from the brown bear (Zou et al., 2022). Interestingly, the FMO3 showed signs of possible pseudogenization in both the polar bear and the sea otter, although it shows similar patterns in other bears and otters. While not a marine mammal, the hippopotamus (*Hippopotamus amphibius*) presents an interesting case, as it is the closest relative of cetaceans that is not fully aquatic but still spends a significant amount of their lives underwater. Furthermore, it is more closely related to cetaceans than any other cetartiodactyl, forming a monophyletic group (Cetancodonta/Whippomorpha) (Zurano et al., 2019). As such, it is interesting to see whether the hippopotamus’ chemical defensome resembles more that of cetaceans or that of other terrestrial cetartiodactyls. We did not observe loss of NR1I2 nor NR1I3 in the hippopotamus, nor did we find the extensive gene loss observed in cetaceans or other marine mammals. This leads us to believe that the reduction of the chemical defensome is cetacean specific, and the shift towards a marine lifestyle had a significant impact on the evolution of the chemical defensome.

Adaptation to the marine environment has significantly influenced the evolution of marine mammals. Multiple studies investigated genomic changes in marine mammals, revealing adaptations to diving, skin morphology, and bone development (Huelsmann et al., 2019; Selleghin-Veiga et al., 2024; Tian et al., 2019; Zhou et al., 2018). In cetaceans, the shift towards the marine environment also led to a change in feeding preference from a predominantly plant-based diet in the ancestor, to a more carnivorous diet today (Thewissen et al., 2009). This shift in diet has more than likely also impacted the genomic gene inventory of cetaceans, multiple genes have been shown to be convergently lost or retained in species with similar diets, including genes of the chemical defensome (Chaney et al., 2026; Hecker et al., 2019; Thomas, 2007; Wagner et al., 2022; Wilhoit et al., 2025). Together, these shifts could have contributed to the observed reduction of the chemical defensome of marine mammals. However, with the increase of marine pollution due to human activity, marine mammals are under considerable chemical stress (Chen et al., 2024; Noël and Brown, 2021). Loss of crucial biotransformation enzymes would mean a significant disadvantage, as it reduced the capacity to detoxify and secrete chemical pollution. In absence of adequate chemical defenses, marine mammals often have no other option than to accumulate these pollutants in their tissue, leading to the high levels of persistent organic pollutants (POPs) in marine wildlife which pose a significant threat to their health (Jepson et al., 2016; Khairy et al., 2021; Vagi et al., 2021; Williams et al., 2023). While many different factors influence the exposure, uptake, and storage of chemical pollution, a recent study of marine mammals in the Barents Sea revealed that the number of functional biotransformation genes in a species’ genome explained the most variation in POP levels in the blubber tissue (Routi et al., in review). This highlights the complexity of the interaction of marine mammals with chemical pollution, and that adaptation to marine life has involved trade-offs in detoxification capacity that may have significant implications for these species’ responses to increasing chemical pollution in marine ecosystems.

## Material & Methods

### Data acquisition

Available annotated reference genomes of all Cetartiodactyla, Carnivora, and Afrotheria were retrieved from NCBI (accessed September 2024) using NCBI datasets (O’Leary et al., 2024). To avoid including highly fragmented genomes, only genomes with a scaffold or contig N50 value of 1Mb or higher were retained. Genomes of known crossbreeds were removed, as were genomes without RefSeq annotations. This resulted in a dataset of 47 Cetartiodactyla (of which 21 Cetacea), 43 Carnivora genomes (of which 9 Pinnipedia), and 7 Afrotheria genomes (of which 1 Sirenia) (Table S2). For each genome annotation, the longest isoform and corresponding protein sequence were extracted for every *locus* using the AGAT toolkit v1.2.0 (Dainat, 2022).

### Identification of defensome genes

An overview of the analysis pipeline can be found in Figure 5. The proteomes of all species, together with the human MANE dataset (Matched Annotation from NCBI and EMBL-EBI; defining one representative protein sequence per locus (Morales et al., 2022)), were divided into orthologous groups using Orthofinder v2.5.5 (Emms and Kelly, 2019). The resulting orthologous groups were functionally annotated by annotating each gene in the group using the eggNOG-mapper v2.1.12 and the eggNOG 5.0.2 database (default parameters, e-value cutoff 1e-10) (Cantalapiedra et al., 2021; Huerta-Cepas et al., 2019). In case of conflicting annotations within the same orthologous group, the most abundant annotation was assigned to the group. Orthologous groups were considered part of the chemical defensome if the annotation of the group matched the names or patterns of defensome genes described by Eide et al. (2021) (available at FAIRDOMHub: https://doi.org/10.15490/FAIRDOMHUB.1.DATAFILE.3957.1). Orthologous groups containing only one gene were excluded from the analysis.

**Figure 5:**
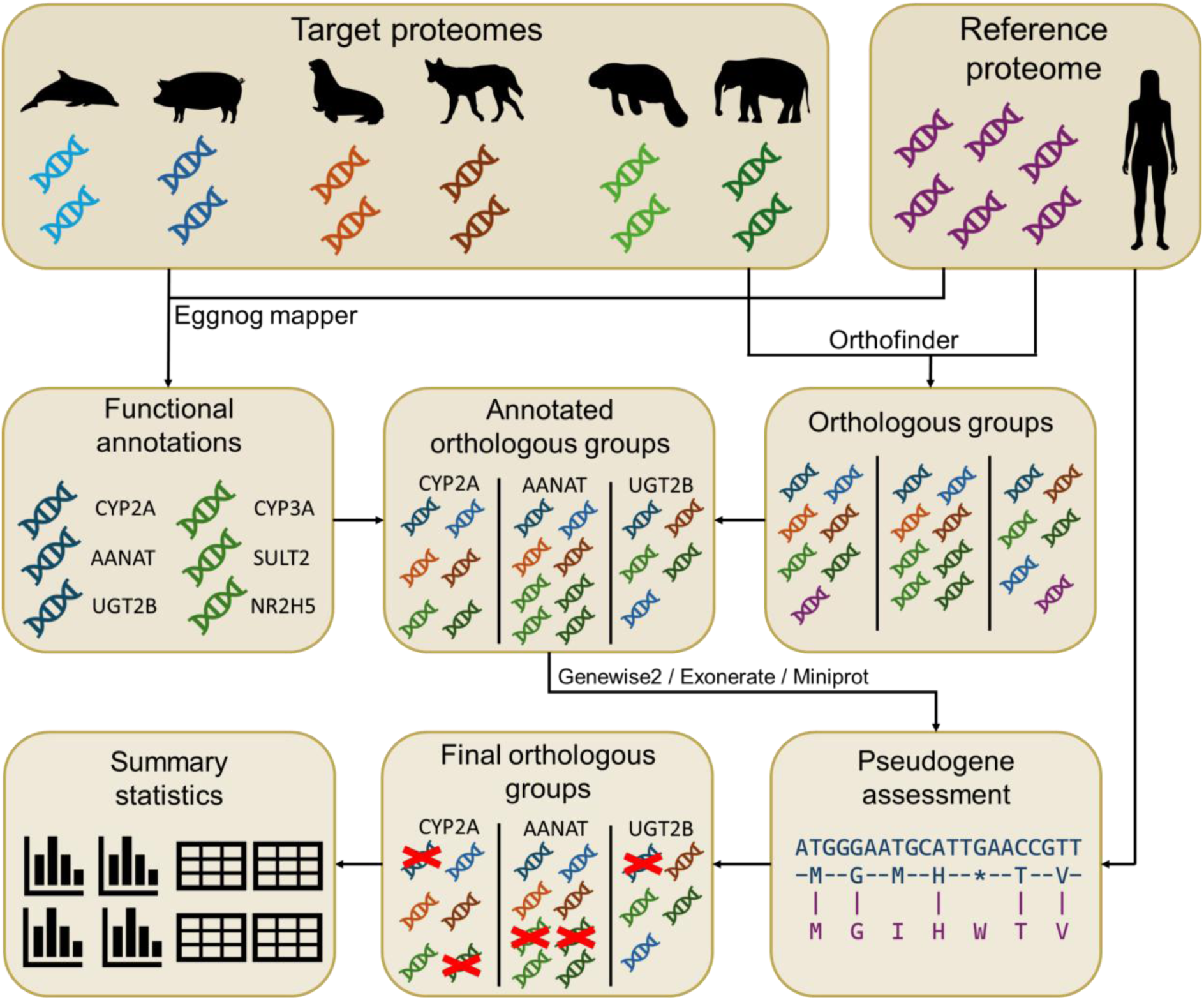
**Graphical overview of the analysis pipeline used to analyse the chemical defensome in the target species.**

### Assessment of pseudogenisation in defensome genes

In orthologous groups that contain at least one human gene, possible pseudogenisation events were investigated. The protein coding sequences of all non-human genes in such orthologous groups were aligned to all human genes in the same group using Diamond (default settings, e-value cutoff 1e-10) (Buchfink et al., 2021). In addition, the protein sequence of the most similar human protein in the orthologous group (as identified by lowest e-value in the Diamond alignments) was aligned to the genomic region (including 25kb up-and downstream) of each non-human gene in the group using GeneWise (v2.4.1, using default parameters, but allowing alignment on both strands (*--both*)) (Birney et al., 2004), Exonerate (v2.4.0, default settings, using the protein to genome model (*--model protein2genome*)) (Slater and Birney, 2005), and Miniprot (v0.15, default settings, but setting the output to alignment (*--aln*)) (Li, 2023). Non-human genes were considered pseudogenes if one or more of the following criteria were met: 1) The protein is shorter than 50% of the protein length of the closest human gene; 2) The protein shows <50% protein identity in the Diamond alignment with the closest human protein; 3) At least one frameshift or premature stop codon was detected in two out of the three protein-to-genome alignments (GeneWise, Exonerate, and Miniprot). In orthologous groups without human reference sequences, all genes were assumed functional.

### Analysis of differences in gene inventory between lineages

Differences in the gene inventory between phylogenetic lineages were assessed in each orthologous group using two different tests. First, for each lineage, the proportions of genomes with at least one functional gene in the orthogroup were compared using Fisher exact tests on the corresponding contingency tables. Second, the average number of functional gene copies in each genome per lineage was compared using non-parametric Mann-Whitney *U* tests. The resulting *p*-values of both tests were adjusted for multiple testing using the Benjamini-Hochberg procedure to control the FDR at 0.05. Statistical analyses were performed using the Pandas and SciPy packages implemented in Python. Lineage-specific differences in gene inventories were manually checked by investigating gene synteny and pseudogene alignments in a selection of species with high assembly continuity (N50).

### Gene synteny of select gene clusters

For a selection of the most variable gene clusters of the chemical defensome (ADH, FMO, CYP2A/B/F/S/G, CYP2C, and UGT2), we constructed synteny maps of representative species within cetaceans (*Orcinus orca*, *Kogia breviceps* and *Balaenoptera musculus)* and pinnipeds (*Mirounga angustirostris*, *Odobenus rosmarus divergens* and *Zalophus californianus)*, as well as reference species, including closely related terrestrials (*Homo sapiens*, *Bos taurus*, *Hippopotamus amphibius*, *Canis lupus familiaris* and *Ursus maritimus*). To build the syntenies, we first identified the neighbouring genes in each defensome gene cluster in *H. sapiens*. We then used these neighbouring genes to extract the sequence of each cluster’s *locus* in all other species. To curate the RefSeq annotations of the defensome genes, we aligned the exons of the *H. sapiens*, *B. taurus* and *C. lupus familiaris* transcripts to their orthologous *loci* using the Geneious R11.1.5 map to reference tool, with parameters set to “high sensitivity”. We then visually inspected these alignments to confirm the sequence of each target gene, taking note of potentially disruptive mutations to flag pseudogenes. Unannotated pseudogenes were also included in the syntenies when more than one third of the exons were located with more than 50% identity with any reference. For labelling of unannotated genes, we used the predicted transcript product in RefSeq’s annotation when applicable, or the name of the cluster otherwise, followed by “-L” for “like”. For the purpose of the main analyses, we treated any gene with disruptive mutations or missing exons as pseudogenes, but a more in-depth breakdown of low-confidence coding sequence disruptions can be found in the supplementary material.

### Gene expression in cetacean liver

We also checked gene expression in each of these clusters to further validate putative pseudogenization using publicly available RNA-sequencing runs for liver tissue. We retrieved runs for *Balaenoptera acutorostrata* (SRR919296), *Delphinus delphis* (ERR1331714, ERR1331719, ERR1331721, SRR22401934), *Delphinapterus leucas* (SRR5282283, SRR5282285, SRR5282288, SRR5282289, SRR5282291, SRR5282292, SRR5282293, SRR5282298), *Eschrichtius robustus* (SRR13853443), *Lagenorhynchus albirostris* (ERR1331697, ERR1331720) and *Monodon monoceros* (SRR8578705) from the European Nucleotide Archive (David et al., 2026). We trimmed the obtained reads to remove any adapters and low-quality bases with Trim Galore v0.6.10, using the default settings for paired end libraries (*--paired*) (Krueger et al., 2023). To filter out rRNA reads we then used SortMeRNA v4.3.7, with the default “smr_v4.3_default_db.fasta” database, in paired-end mode and keeping the read pair if at least one of them did not align (*--fastx --out2 --paired_out --other*) (Kopylova et al., 2012). We mapped the filtered reads to their respective reference genome using STAR v2.7.11b, with the parameters *--outFilterMultimapNmax 20 --outFilterMismatchNmax 999 --outFilterMismatchNoverReadLmax 0.04 –alignIntronMax 1000000 --alignMatesGapMax 1000000 --peOverlapNbasesMin 20*. The genomes were indexed with default parameters. Finally, we computed coverage from these alignments using bamCoverage v3.5.6 (Ramírez et al., 2016) with default settings, for use with pyGenomeTracks v3.9 (Lopez-Delisle et al., 2021) to produce coverage figures.

## Data availability

The analysis pipeline implemented in Snakemake is available on Github (https://github.com/bramdanneels/MarineMammalDefensome). All chemical defensome data (per-gene information regarding annotation, orthogroup membership, and pseudogene assessment; per-orthogroup information regarding gene content; per-orthogroup comparisons between phylogenetic lineages) is available on Zenodo (https://zenodo.org/records/19920065).

## Acknowledgements

This work was supported by the Norwegian Research Council (NFR) grants 334739 (Marmadetox) and 326819 (EBP-Nor) and by the Foundation for Science and Technology (FCT), Portugal: research grant awarded to D.O. (https://doi.org/10.54499/2021.06293.BD), 2023.07615.CEECIND/CP2848/CT0007 awarded to R.R. and by strategic funding awarded to CIIMAR (UIDB/04423/2020 and UIDP/04423/2020).

## Notes

https://zenodo.org/records/19920065

https://github.com/bramdanneels/MarineMammalDefensome

### Competing Interest Statement

The authors have declared no competing interest.

