## Supplementary Figures S1-S11 for "Convergent gene erosion in the chemical defensome of marine mammals"

**Figure S1: Gene count and distribution of reductases and various stress responses of the chemical defensome of Cetartiodactyla, Carnivora, and Afrothera.** Colours on the left correspond to the different investigated lineages. Dark green: Afrotheria (without Sirenia); light green: Sirenia; brown: Carnivora (without Pinnipedia); orange: Pinnipedia; dark blue: Cetartiodactyla (without Cetacea); light blue: Cetacea. Heatmap colours correspond to the number of detected functional genes in each gene family.

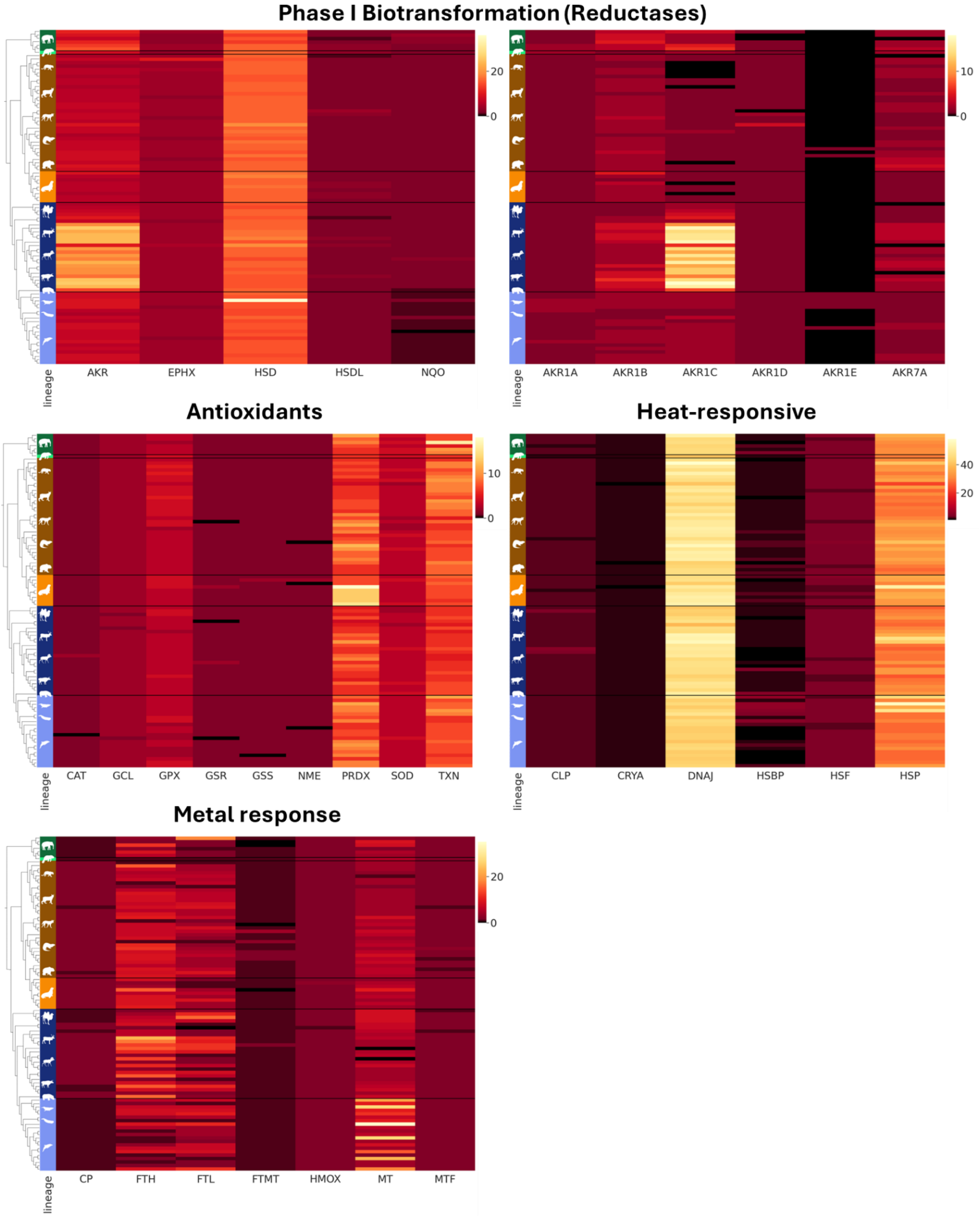

**Figure S2: Synteny of the ADH gene cluster in Cetartiodactyla and Carnivora.** Schematic representation of the aldehyde dehydrogenase gene cluster in select species, including humans; Cetartiodactyla, including a representative for terrestrial and semi-aquatic members, as well as for Odontoceti, Physeteroidea and Mysticeti; and Carnivora, including a representative for terrestrial and semi-aquatic members, as well as for Phocidae, Odobenidae and Otariidae. Neighbouring genes are shown in gray, ADH genes in yellow, and ADH pseudogenes in stripes. Specifically here, for the protein product of *Odobenus rosmarus divergens*, ADH1C was considered pseudogenized due to a single nucleotide deletion at the very beginning of exon 5; however, it is possible that non-canonical splice sites that do not disrupt the reading frame exist in this exon or, since it’s a single mutation, it could be a non-ubiquitous polymorphism. The *Kogia breviceps* silhouette was adapted from its depiction by “Chris huh”, under the [CC BY 3.0 license](https://creativecommons.org/licenses/by/3.0/).

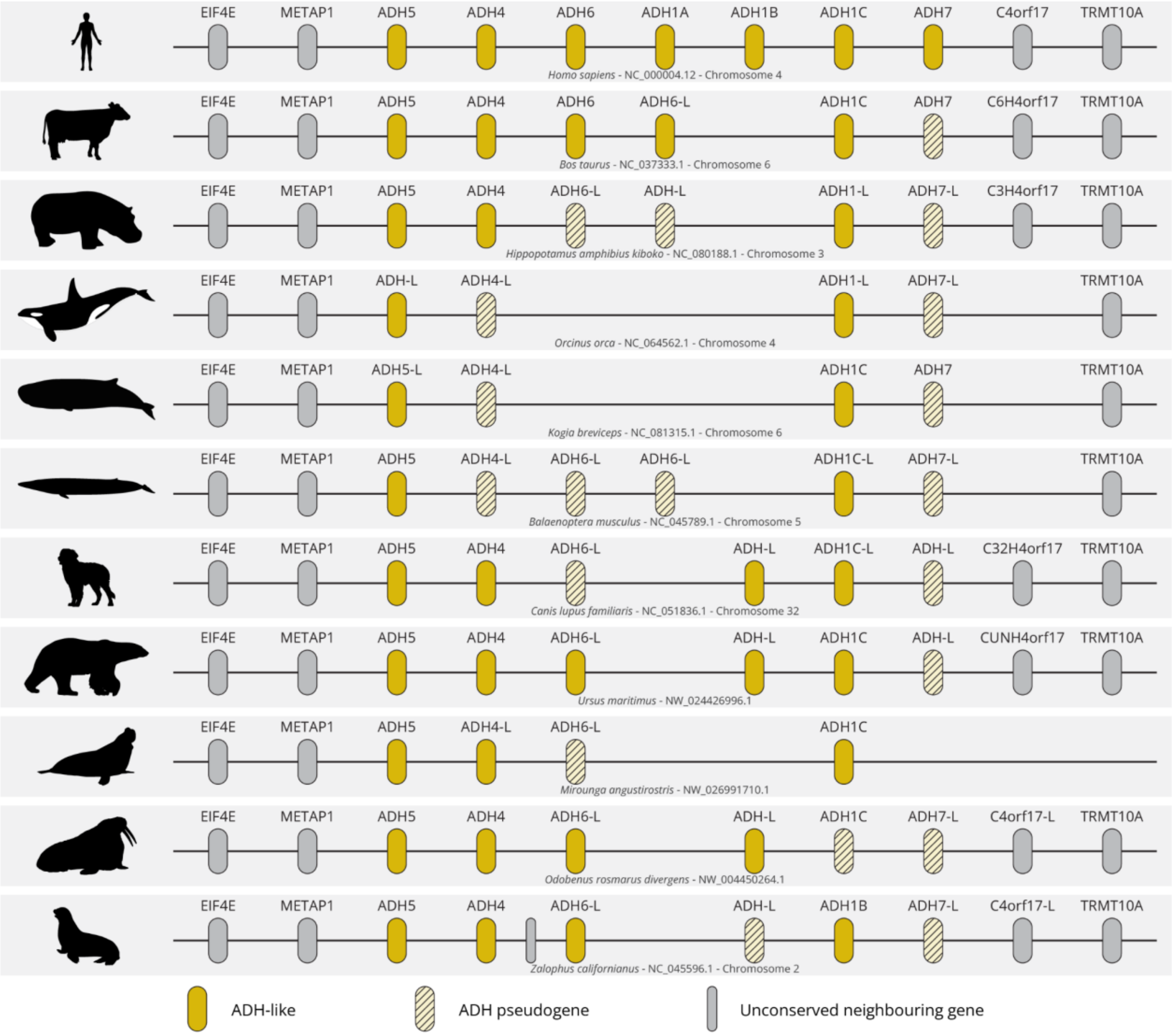


**Figure S3: Gene expression in the ADH cluster in cetacean liver tissue.** Gene expression coverage for the analysed cetaceans for the ADH gene cluster, including a portion of the flanking genes. For each species, the top track graphs the log1p value of raw reads per bin, and the bottom track the longest transcript of each gene, in the specified genomic region. Gene names were included for ADH transcripts. For *Delphinus delphis*, *Delphinapterus leucas* and *Lagenorhynchus albirostris*, the coverage tracks overlay the results for a total of four, eight, and two sequencing runs, respectively.

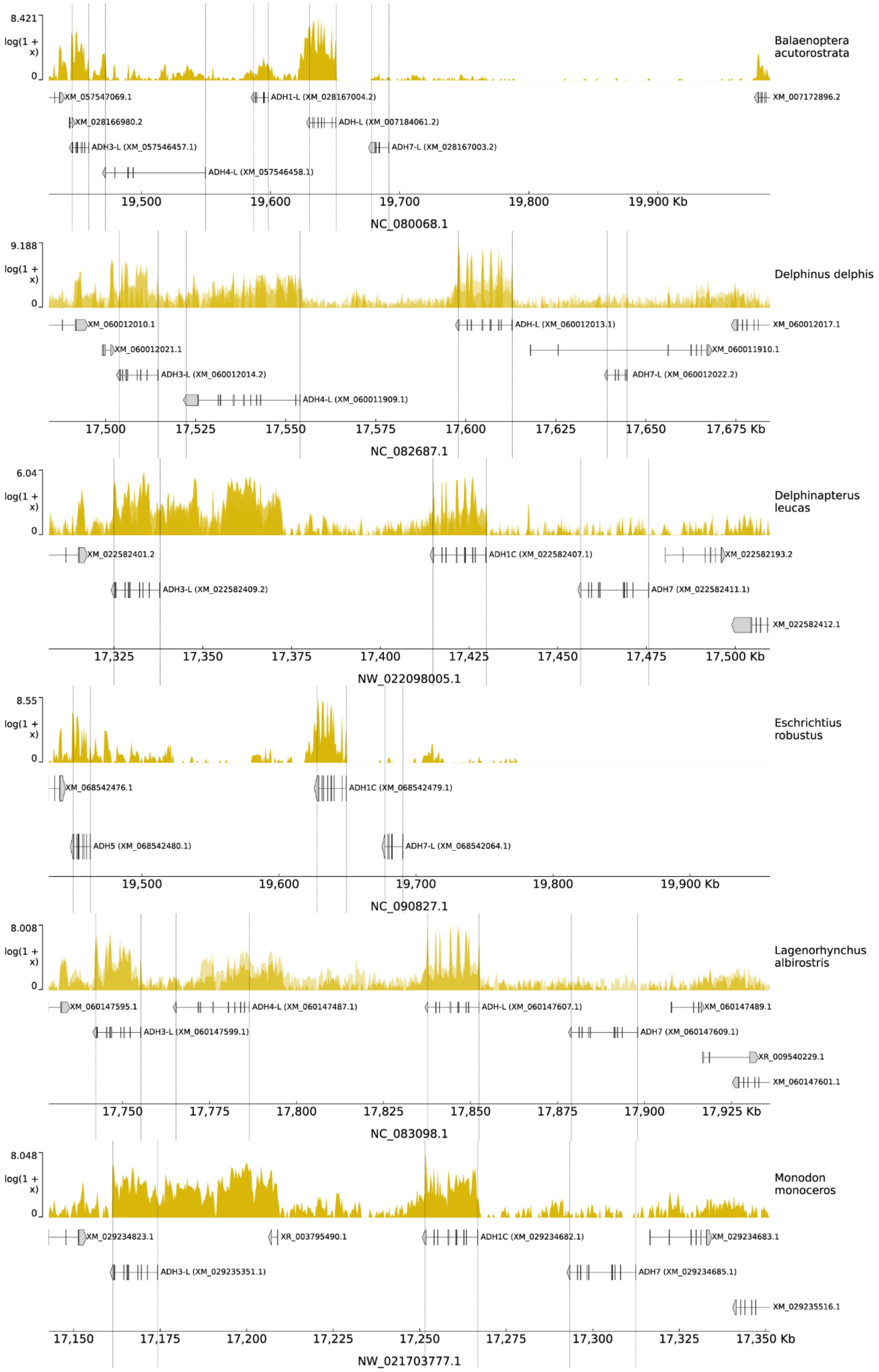
8.421
log(1 + x)
0
Balaenoptera acutorostrata
XM_057547069.1
ADH1-L (XM_028167004.2)
XM_007172896.2
XM_028166980.2
ADH-L (XM_007184061.2)
ADH3-L (XM_057546457.1)
ADH7-L (XM_028167003.2)
ADH4-L (XM_057546458.1)
19,500
19,600
19,700
19,800
19,900 Kb
NC_080068.1
9.188
log(1 + x)
0
Delphinus delphis
XM_060012010.1
ADH-L (XM_060012013.1)
XM_060012017.1
XM_060012021.1
XM_060011910.1
ADH3-L (XM_060012014.2)
ADH7-L (XM_060012022.2)
ADH4-L (XM_060011909.1)
17,500
17,525
17,550
17,575
17,600
17,625
17,650
17,675 Kb
NC_082687.1
6.04
log(1 + x)
0
Delphinapterus leucas
XM_022582401.2
ADH1C (XM_022582407.1)
XM_022582193.2
ADH3-L (XM_022582409.2)
ADH7 (XM_022582411.1)
XM_022582412.1
17,325
17,350
17,375
17,400
17,425
17,450
17,475
17,500 Kb
NW_022098005.1
8.55
log(1 + x)
0
Eschrichtius robustus
XM_068542476.1
ADH1C (XM_068542479.1)
ADH5 (XM_068542480.1)
ADH7-L (XM_068542064.1)
19,500
19,600
19,700
19,800
19,900 Kb
NC_090827.1
8.008
log(1 + x)
0
Lagenorhynchus albirostris
XM_060147595.1
ADH4-L (XM_060147487.1)
ADH-L (XM_060147607.1)
XM_060147489.1
ADH3-L (XM_060147599.1)
ADH7 (XM_060147609.1)
XR_009540229.1
XM_060147601.1
17,750
17,775
17,800
17,825
17,850
17,875
17,900
17,925 Kb
NC_083098.1
8.048
log(1 + x)
0
Monodon monoceros
XM_029234823.1
XR_003795490.1
ADH1C (XM_029234682.1)
XM_029234683.1
ADH3-L (XM_029235351.1)
ADH7 (XM_029234685.1)
XM_029235516.1
17,150
17,175
17,200
17,225
17,250
17,275
17,300
17,325
17,350 Kb
NW_021703777.1

**Figure S4: Synteny of the FMO gene cluster in Cetartiodactyla and Carnivora.** Schematic representation of the flavin-containing dehydrogenase gene cluster in select species, including humans; Cetartiodactyla, including a representative for terrestrial and semi-aquatic members, as well as for Odontoceti, Physeteroidea and Mysticeti; and Carnivora, including a representative for terrestrial and semi-aquatic members, as well as for Phocidae, Odobenidae and Otariidae. In mammals, this cluster does not include FMO5. Neighbouring genes are shown in gray, FMO genes in red, and FMO pseudogenes in stripes. Of note, there's a large stretch of 16 million nucleotides splitting the cluster in *Kogia breviceps*, likely due to genomic rearrangements. Regarding specific genes, *Ursus maritimus*' FMO3 has 2 disruptive mutations, both found at the end of exon 5. However, the possibility of alternative splicing that skips both mutations cannot be discarded. The FMO1 of *Mirounga angustirostris* contains a single non-silent, nonsense mutation, in the middle of exon 5. Thus, alternative splicing that does not terminate the reading frame is possible or, since it's a single mutation across the whole gene, it could be a non-ubiquitous polymorphism. For *Odobenus rosmarus divergens*, the retrieved FMO1 sequence is partial, encompassing only exons 2 to 4 out of a total of 8. There are neighbouring assembly gaps capable of hiding the remaining exons, so the gene was treated as coding. The *Kogia breviceps* silhouette was adapted from its depiction by "Chris huh", under the [CC BY 3.0 license](https://creativecommons.org/licenses/by/3.0/).

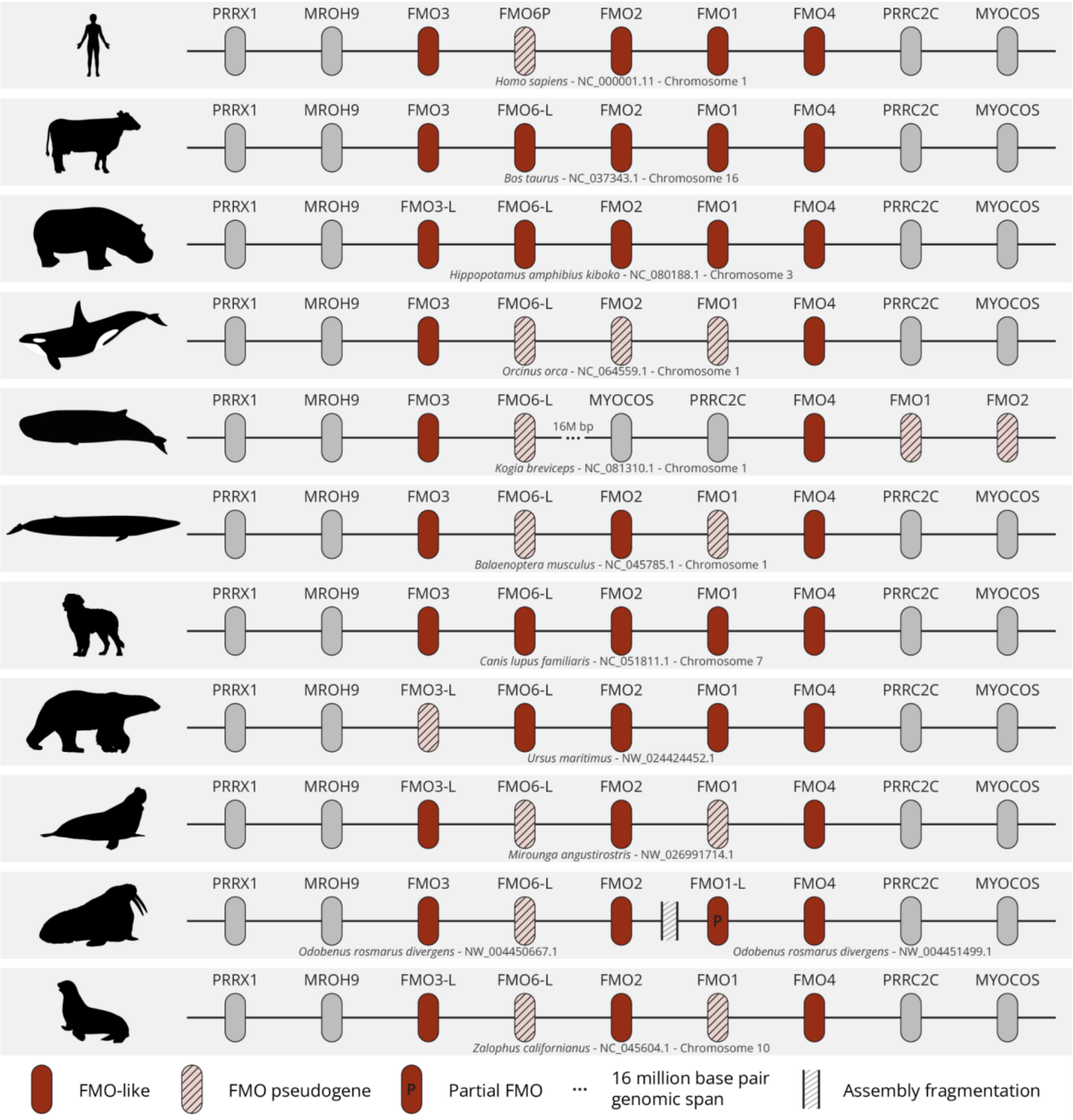

**Figure S5: Gene expression in the FMO cluster in cetacean liver tissue.** Gene expression coverage for the analysed cetaceans for the FMO gene cluster, including a portion of the flanking genes. For each species, the top track graphs the log1p value of raw reads per bin, and the bottom track the longest transcript of each gene, in the specified genomic region. Gene names were included for FMO transcripts. For *Delphinus delphis*, *Delphinapterus leucas* and *Lagenorhynchus albirostris*, the coverage tracks overlay the results for a total of four, eight, and two sequencing runs, respectively.

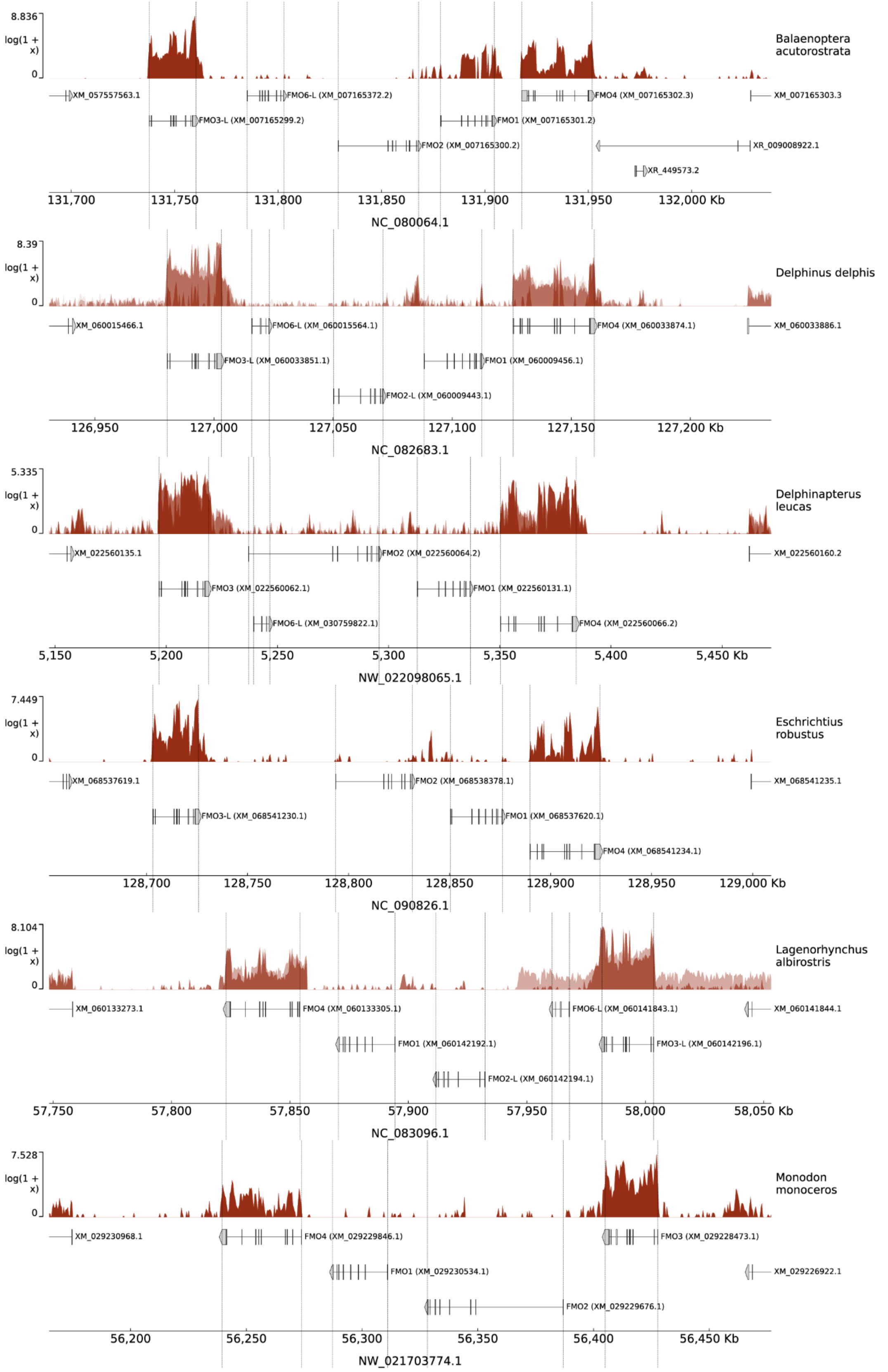

**Figure S6: Synteny of the CYP2A/B/F/G/S gene cluster in Cetartiodactyla and Carnivora.** Schematic representation of the cytochrome P450 2A/B/F/G/S gene cluster in select species, including humans; Cetartiodactyla, including a representative for terrestrial and semi-aquatic members, as well as for Odontoceti, Physeteroidea and Mysticeti; and Carnivora, including a representative for terrestrial and semi-aquatic members, as well as for Phocidae, Odobenidae and Otariidae. Neighbouring genes are shown in gray, CYP genes in green, and CYP pseudogenes in stripes. Specifically, *Canis lupus familiaris*' CYP2S1 has a single disruptive mutation, a deletion at the end of exon 5. However, since it is a single mutation, it could be a non-ubiquitous polymorphism. In fact, this mutation is not present in a different breed's assembly, the German Shepherd's (located in chromosome 1 with the accession number NC_049222.1). The *Kogia breviceps* silhouette was adapted from its depiction by "Chris huh", under the CC BY 3.0 license.

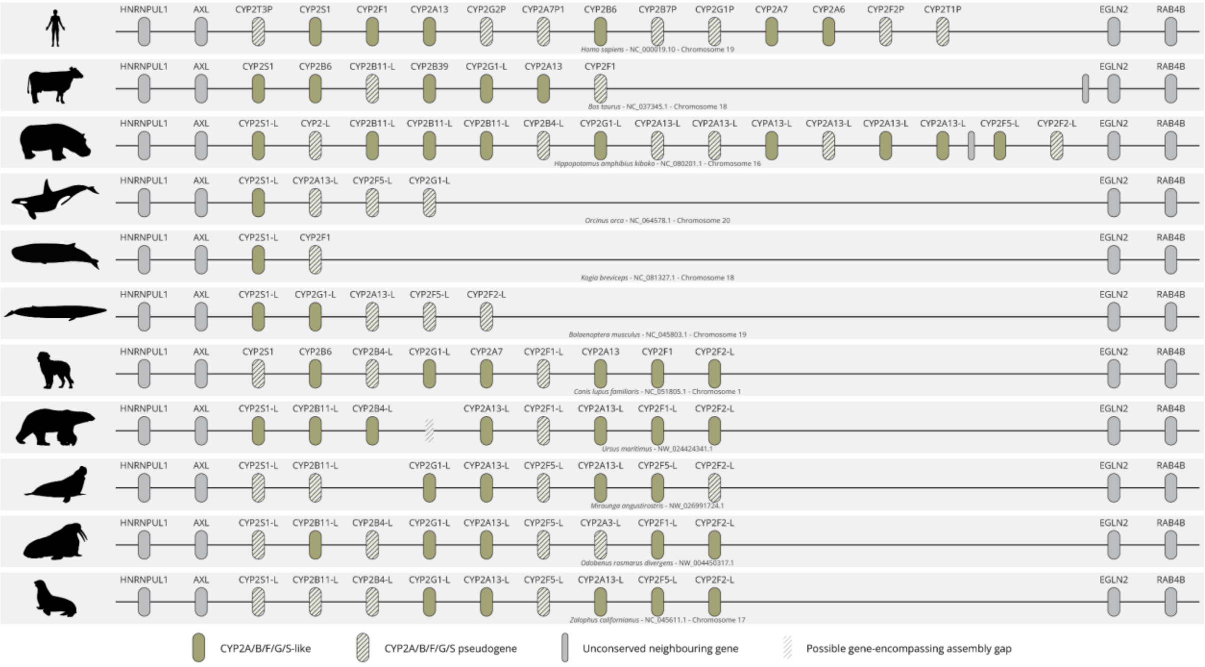


**Figure S7: Gene expression in the CYP2A/B/F/G/S cluster in cetacean liver tissue.** Gene expression coverage for the analysed cetaceans for the CYP2A/B/F/G/S gene cluster, including a portion of the flanking genes. For each species, the top track graphs the log1p value of raw reads per bin, and the bottom track the longest transcript of each gene, in the specified genomic region. Gene names were included for CYP family transcripts. For *Delphinus delphis*, *Delphinapterus leucas* and *Lagenorhynchus albirostris*, the coverage tracks overlay the results for a total of four, eight, and two sequencing runs, respectively.

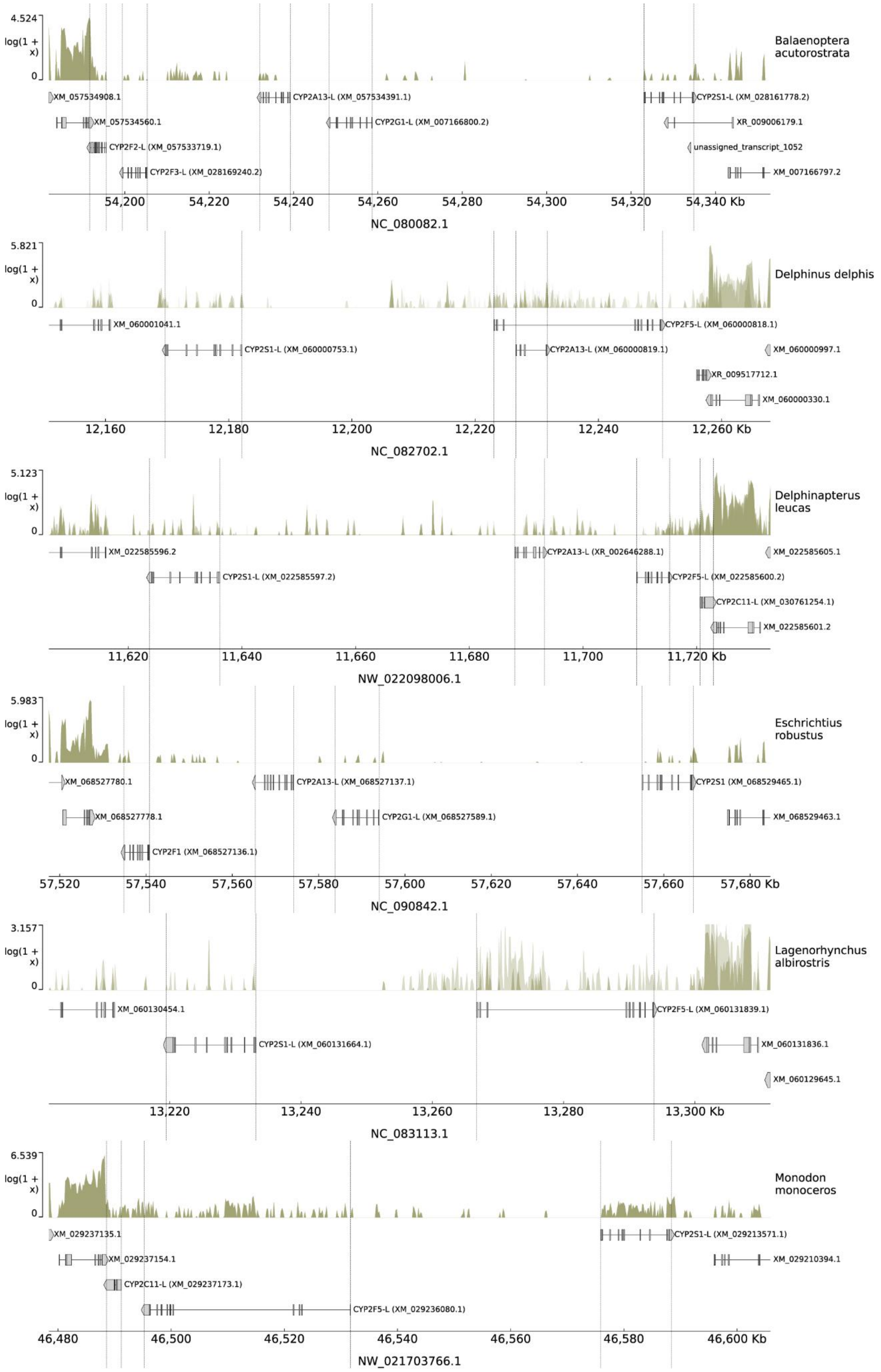

4.524
log(1 + x)
0
Balaenoptera acutorostrata
XM_057534908.1
CYP2A13-L (XM_057534391.1)
CYP2S1-L (XM_028161778.2)
XM_057534560.1
CYP2G1-L (XM_007166800.2)
XR_009006179.1
CYP2F2-L (XM_057533719.1)
unassigned_transcript_1052
CYP2F3-L (XM_028169240.2)
XM_007166797.2
54,200
54,220
54,240
54,260
54,280
54,300
54,320
54,340 Kb
NC_080082.1
5.821
log(1 + x)
0
Delphinus delphis
XM_060001041.1
CYP2F5-L (XM_060000818.1)
CYP2S1-L (XM_060000753.1)
CYP2A13-L (XM_060000819.1)
XM_060000997.1
XR_009517712.1
XM_060000330.1
12,160
12,180
12,200
12,220
12,240
12,260 Kb
NC_082702.1
5.123
log(1 + x)
0
Delphinapterus leucas
XM_022585596.2
CYP2A13-L (XR_002646288.1)
XM_022585605.1
CYP2S1-L (XM_022585597.2)
CYP2F5-L (XM_022585600.2)
CYP2C11-L (XM_030761254.1)
XM_022585601.2
11,620
11,640
11,660
11,680
11,700
11,720 Kb
NW_022098006.1
5.983
log(1 + x)
0
Eschrichtius robustus
XM_068527780.1
CYP2A13-L (XM_068527137.1)
CYP2S1 (XM_068529465.1)
XM_068527778.1
CYP2G1-L (XM_068527589.1)
XM_068529463.1
CYP2F1 (XM_068527136.1)
57,520
57,540
57,560
57,580
57,600
57,620
57,640
57,660
57,680 Kb
NC_090842.1
3.157
log(1 + x)
0
Lagenorhynchus albirostris
XM_060130454.1
CYP2F5-L (XM_060131839.1)
CYP2S1-L (XM_060131664.1)
XM_060131836.1
XM_060129645.1
13,220
13,240
13,260
13,280
13,300 Kb
NC_083113.1
6.539
log(1 + x)
0
Monodon monoceros
XM_029237135.1
CYP2S1-L (XM_029213571.1)
XM_029237154.1
XM_029210394.1
CYP2C11-L (XM_029237173.1)
CYP2F5-L (XM_029236080.1)
46,480
46,500
46,520
46,540
46,560
46,580
46,600 Kb
NW_021703766.1

**Figure S8: Synteny of the CYP2C gene cluster in Cetartiodactyla and Carnivora.** Schematic representation of the cytochrome P450 2C gene cluster in select species, including humans; Cetartiodactyla, including a representative for terrestrial and semi-aquatic members, as well as for Odontoceti, Physeteroidea and Mysticeti; and Carnivora, including a representative for terrestrial and semi-aquatic members, as well as for Phocidae, Odobenidae and Otariidae. Neighbouring genes are shown in gray, CYP2C genes in green, and CYP2C pseudogenes in stripes. Here, while the *Canis lupus familiaris* synteny for the RefSeq reference genome has only one CYP2C gene, a different assembly for the Basenji breed has two coding genes (located in chromosome 28 with the accession number NC_049769.1). One of the genes for *Ursus maritimus* is shown as a pseudogene since it is missing the first 3 exons. However, the assembly has gaps that could be hiding additional CYP2C genes which exist in the closely related *Ursus arctos'* genome. In fact, its CYP2C in the same position as *Ursus maritimus'* pseudogene is missing the same 3 exons, and the annotation includes it as part of the other CYP2C; furthermore, the DNA strand the genes are in in both species is coherent with the hypothesis that the *Ursus maritimus* assembly does not exhaustively portray this *locus* in the species (not shown). The *Kogia breviceps* silhouette was adapted from its depiction by “Chris huh”, under the CC BY 3.0 license.

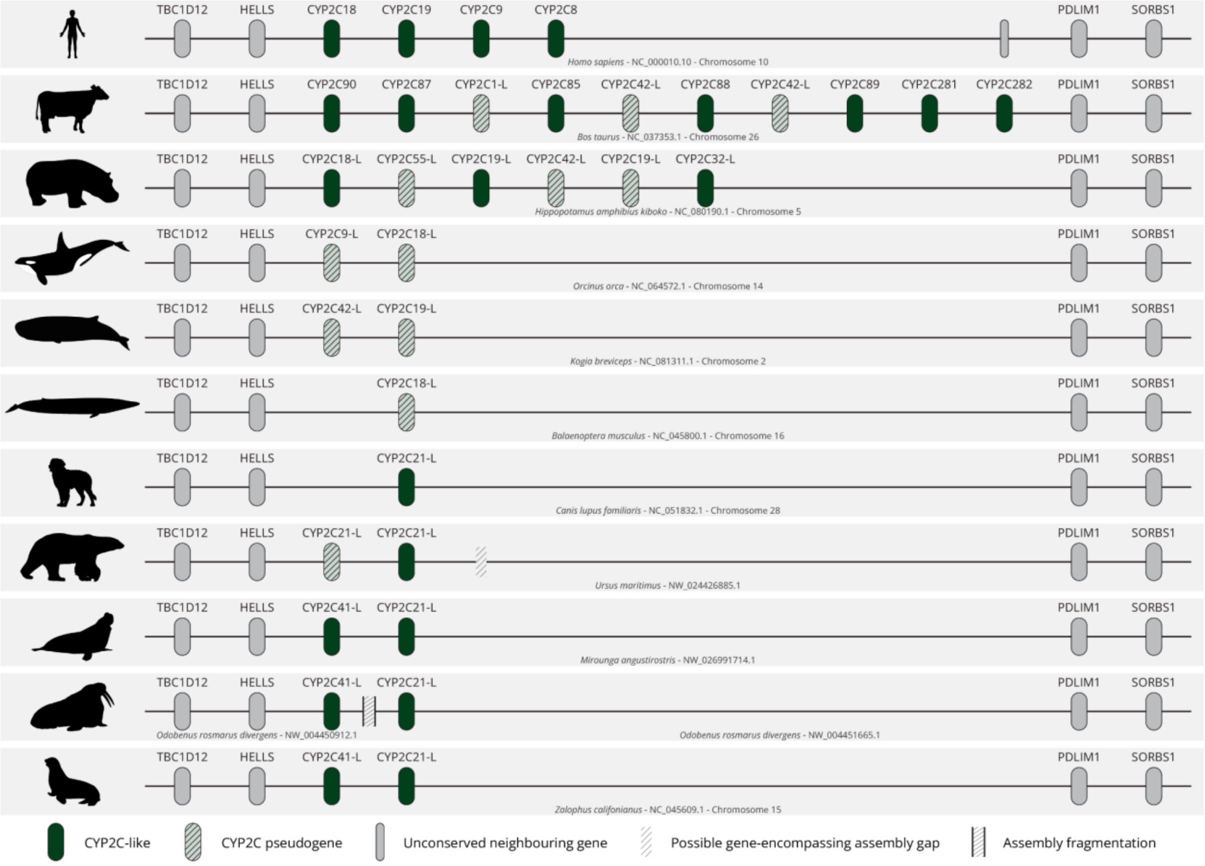

**Figure S9: Gene expression in the CYP2C cluster in cetacean liver tissue.** Gene expression coverage for the analysed cetaceans for the CYP2C gene cluster, including a portion of the flanking genes. For each species, the top track graphs the log1p value of raw reads per bin, and the bottom track the longest transcript of each gene, in the specified genomic region. Gene names were included for CYP family transcripts. For *Delphinus delphis*, *Delphinapterus leucas* and *Lagenorhynchus albirostris*, the coverage tracks overlay the results for a total of four, eight, and two sequencing runs, respectively.

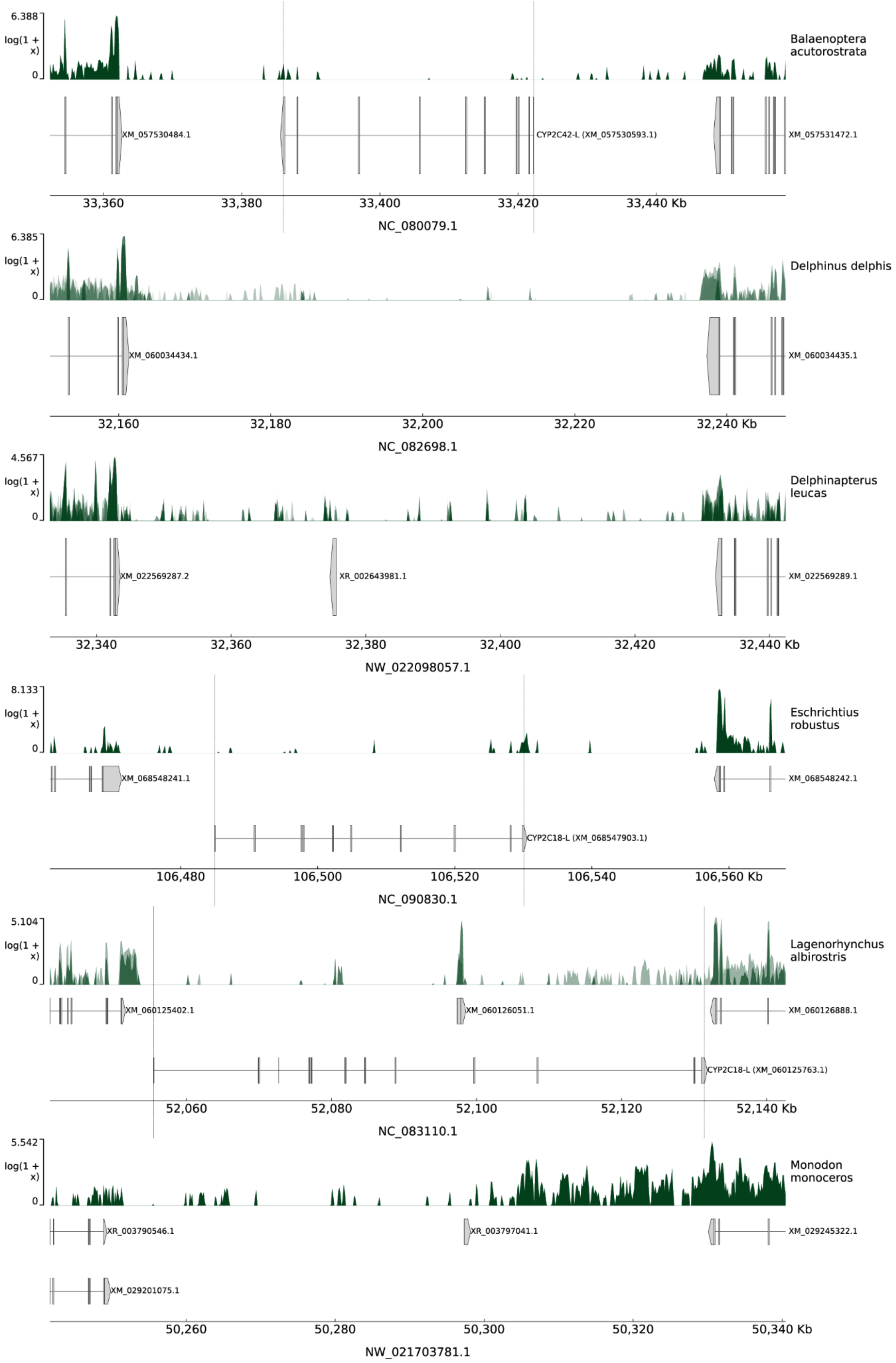

**Figure S10: Synteny of the UGT2 gene cluster in Cetartiodactyla and Carnivora.** Schematic representation of the UDP-glucuronosyltransferase 2 gene cluster in select species, including humans; Cetartiodactyla, including a representative for terrestrial and semi-aquatic members, as well as for Odontoceti, Physeteroidea and Mysticeti; and Carnivora, including a representative for terrestrial and semi-aquatic members, as well as for Phocidae, Odobenidae and Otariidae. Neighbouring genes are shown in gray, UGT2 genes in green, and UGT2 pseudogenes in stripes. Human UGT2A1 and UGT2A2, historically treated as two separate genes, are known to be exon-sharing transcripts of the same gene. They use alternative starting exons, but the remaining 5 exons are the same. However, most of the functionally relevant part of the sequence is included in the first exon. Thus, in the scope of this synteny, each transcript was treated as an individual gene, and each starting exon is represented inside a larger box. Human pseudogenes in this cluster are very numerous, so they were omitted. Regarding specific genes, *Orcinus orca*'s UGT2A1-L was considered a pseudogene because exon 2 was not found. However, one copy of the highly relevant exon 1 is intact. Thus, the possibility that the gene makes use of an alternative that is coding cannot be ruled out. The *Kogia breviceps* silhouette was adapted from its depiction by "Chris huh", under the CC BY 3.0 license.

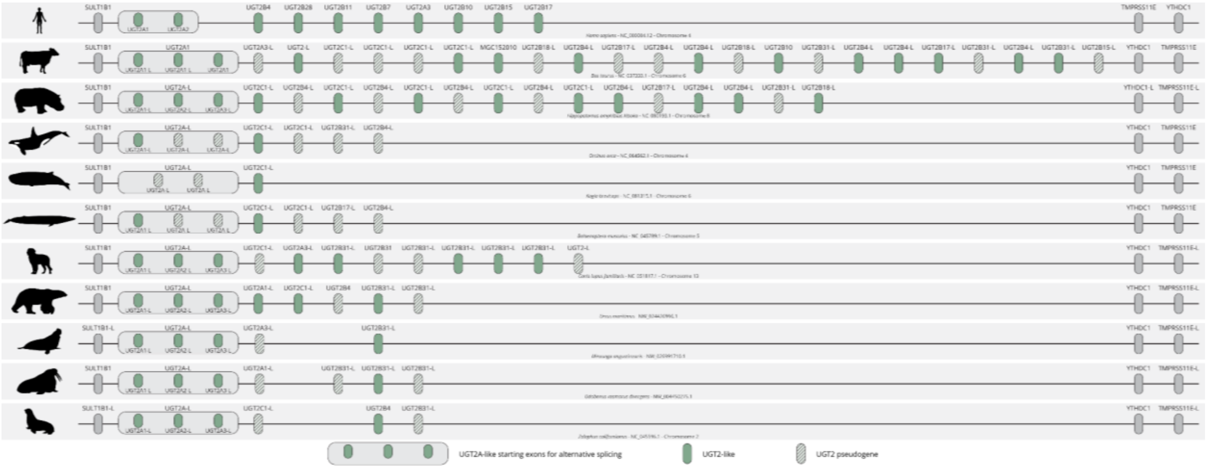

**Figure S11: Gene expression in the UGT2 cluster in cetacean liver tissue.** Gene expression coverage for the analysed cetaceans for the UGT2 gene cluster, including a portion of the flanking genes. For each species, the top track graphs the log1p value of raw reads per bin, and the bottom track the longest transcript of each gene, in the specified genomic region. Gene names were included for UGT transcripts. For *Delphinus delphis*, *Delphinapterus leucas* and *Lagenorhynchus albirostris*, the coverage tracks overlay the results for a total of four, eight, and two sequencing runs, respectively.

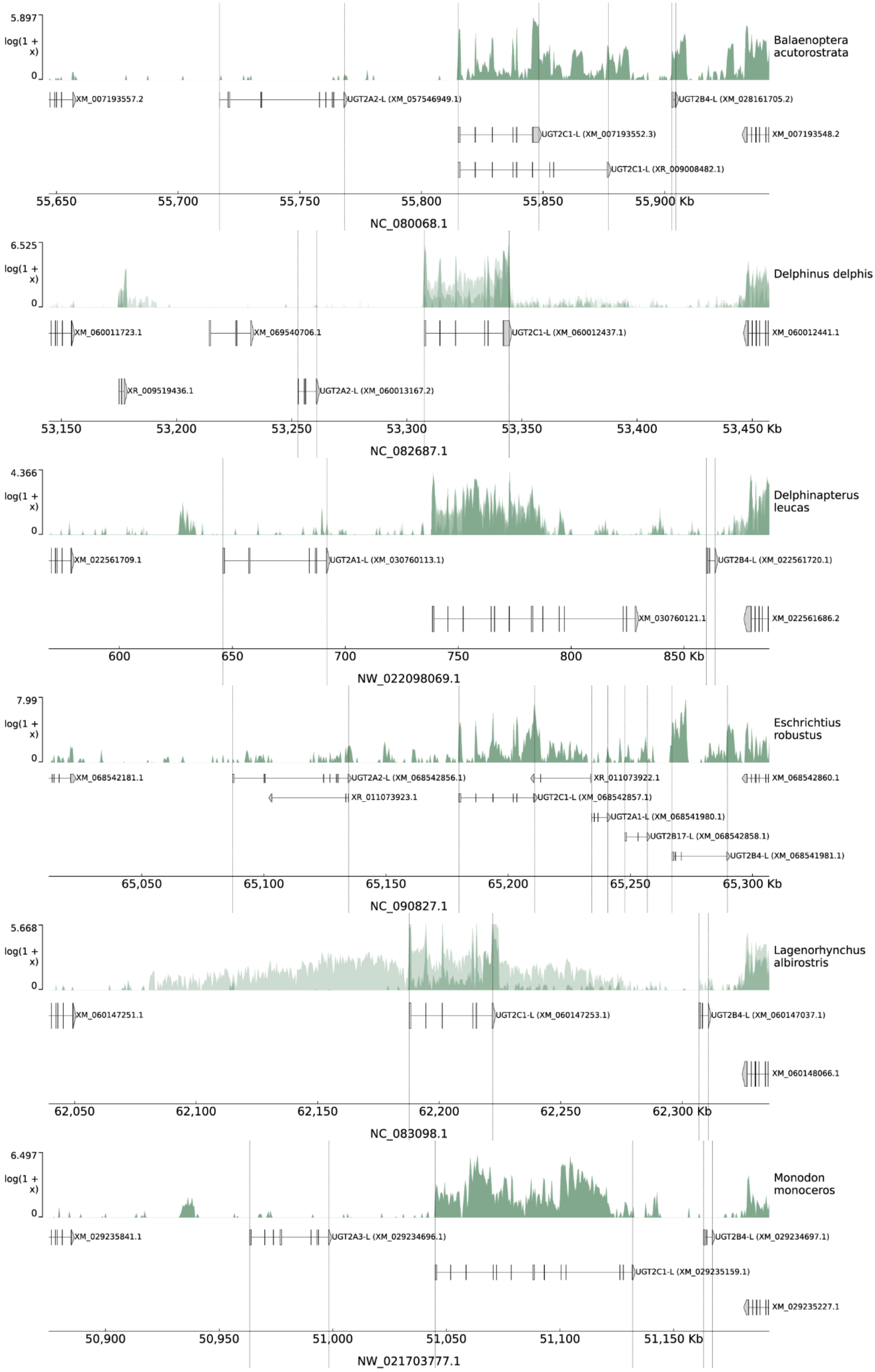
5.897
log(1 + x)
0
Balaenoptera acutorostrata
XM_007193557.2
UGT2A2-L (XM_057546949.1)
UGT2B4-L (XM_028161705.2)
UGT2C1-L (XM_007193552.3)
XM_007193548.2
UGT2C1-L (XR_009008482.1)
55,650
55,700
55,750
55,800
55,850
55,900 Kb
NC_080068.1
6.525
log(1 + x)
0
Delphinus delphis
XM_060011723.1
XM_069540706.1
UGT2C1-L (XM_060012437.1)
XM_060012441.1
XR_009519436.1
UGT2A2-L (XM_060013167.2)
53,150
53,200
53,250
53,300
53,350
53,400
53,450 Kb
NC_082687.1
4.366
log(1 + x)
0
Delphinapterus leucas
XM_022561709.1
UGT2A1-L (XM_030760113.1)
UGT2B4-L (XM_022561720.1)
XM_030760121.1
XM_022561686.2
600
650
700
750
800
850 Kb
NW_022098069.1
7.99
log(1 + x)
0
Eschrichtius robustus
XM_068542181.1
UGT2A2-L (XM_068542856.1)
XR_011073922.1
XM_068542860.1
XR_011073923.1
UGT2C1-L (XM_068542857.1)
UGT2A1-L (XM_068541980.1)
UGT2B17-L (XM_068542858.1)
UGT2B4-L (XM_068541981.1)
65,050
65,100
65,150
65,200
65,250
65,300 Kb
NC_090827.1
5.668
log(1 + x)
0
Lagenorhynchus albirostris
XM_060147251.1
UGT2C1-L (XM_060147253.1)
UGT2B4-L (XM_060147037.1)
XM_060148066.1
62,050
62,100
62,150
62,200
62,250
62,300 Kb
NC_083098.1
6.497
log(1 + x)
0
Monodon monoceros
XM_029235841.1
UGT2A3-L (XM_029234696.1)
UGT2B4-L (XM_029234697.1)
UGT2C1-L (XM_029235159.1)
XM_029235227.1
50,900
50,950
51,000
51,050
51,100
51,150 Kb
NW_021703777.1
